# The molecular basis for DNA-binding by competence T4P is distinct in Gram-positive and Gram-negative species

**DOI:** 10.1101/2025.02.17.638644

**Authors:** Nicholas D. Christman, Ankur B. Dalia

**Affiliations:** Department of Biology, Indiana University, Bloomington, IN

## Abstract

Competence type IV pili (T4P) are bacterial surface appendages that facilitate DNA uptake during horizontal gene transfer by natural transformation. These dynamic structures actively extend from the cell surface, bind to DNA in the environment, and then retract to import bound DNA into the cell. Competence T4P are found in diverse Gram-negative (diderm) and Gram-positive (monoderm) bacterial species. While the mechanism of DNA-binding by diderm competence T4P has been the recent focus of intensive study, relatively little is known about DNA-binding by monoderm competence T4P. Here, we use *Streptococcus pneumoniae* as a model system to address this question. Competence T4P likely bind to DNA via a tip-associated complex of proteins called minor pilins, and recent work highlights a high degree of structural conservation between the minor pilin tip complexes of monoderm and diderm competence T4P. In diderms, positively charged residues in one minor pilin, FimT, are critical for DNA-binding. We show that while these residues are conserved in ComGD, the FimT homolog of monoderms, they only play a minor role in DNA uptake for natural transformation. Instead, we find that two-positively charged residues in the neighboring minor pilin, ComGF (the PilW homolog of monoderms), play the dominant role in DNA uptake for natural transformation. Furthermore, we find that these residues are conserved in other monoderms, but not diderms. Together, these results suggest that the molecular basis for DNA-binding has either diverged or evolved independently in monoderm and diderm competence T4P.

**AUTHOR SUMMARY:** Diverse bacteria use extracellular structures called competence type IV pili (T4P) to take up DNA from their environment. The uptake of DNA by T4P is the first step of natural transformation, a mode of horizontal gene transfer that contributes to the spread of antibiotic resistance and virulence traits in diverse clinically relevant Gram-negative (diderm) and Gram-positive (monoderm) bacterial species. While the mechanism of DNA binding by competence T4P in diderms has been an area of recent study, relatively little is known about how monoderm competence T4P bind DNA. Here, we explore how monoderm competence T4P bind DNA using *Streptococcus pneumoniae* as a model system. Our results indicate that while monoderm T4P and diderm T4P likely have conserved structural features, the DNA-binding mechanism of each system is distinct.

## INTRODUCTION

Natural transformation (NT; also known as genetic transformation or natural competence) is a broadly conserved mechanism of horizontal gene transfer in diverse bacteria and archaea [1]. During this process, cells take up free DNA from the environment to integrate into their genome by homologous recombination. The first step of NT is the uptake of extracellular DNA, which is facilitated by dynamic surface appendages called competence T4P.

Competence T4P actively extend into the extracellular environment, bind to free DNA, and then retract to facilitate the uptake of DNA, as shown in both the diderm *Vibrio cholerae* [2] and the monoderm *S. pneumoniae* [3]. The active extension and retraction of pili is supported by an envelope-spanning molecular machine that is powered by cytoplasmic ATPase motors [4–6]. Through this activity, competence T4P facilitate the uptake of a bight of double-stranded DNA into the periplasm in diderms, or the space between the cell wall and cytoplasmic membrane in monoderms (i.e., the “Gram-positive periplasm” [7]). This bight of DNA is then bound by ComEA, a periplasmic (diderm) or membrane-embedded (monoderm) DNA-binding protein, which functions as a molecular ratchet to further drive DNA uptake [8–10]. In both diderms and monoderms, a single strand of this DNA is then translocated across the cytoplasmic membrane where it can then be integrated into the host genome by homologous recombination.

In T4P, a group of proteins called minor pilins form a complex that initiates the assembly of the pilus filament; thus, this minor pilin complex is located at the tip of extended pili [11–13]. Tip-associated DNA-binding has been observed in both monoderms and diderms [2, 3]. DNA-binding by competence T4P can occur in either a sequence-dependent or sequence-independent manner [14, 15], however most naturally competent organisms exhibit sequence-independent DNA uptake. Diverse proteins that bind DNA in a sequence-independent manner rely on charge-based interactions [16–18]. Altogether, this supports the hypothesis that competence T4P bind DNA via an electrostatic interaction between positively charged residues in the minor pilins and the negatively charged phosphate backbone of DNA. Consistent with this, recent work from a number of groups demonstrates that positively charged residues at the C-terminus of the minor pilin FimT contribute to the DNA-binding activity of competence T4P in a number of diderms, including *V. cholerae* [2], *Legionella pneumophila* [19], *Xylella fastidiosa* [20], and *Acinetobacter baylyi* [21]. The molecular basis for DNA-binding by monoderm competence T4P, however, remains poorly understood. Recent work highlights that the overall structure of the minor pilin tip complex of the *Streptococcus sanguinis* competence T4P is highly similar to the tip complex of diderm T4P and type 2 secretion systems (T2SSs) [6, 22–24]. This suggests that the molecular basis for DNA-binding may be conserved across diderms and monoderms. In this study, we sought to directly test this hypothesis.

## RESULTS

### Positively charged residues at the C-terminus of FimT_Vc_ are critical for DNA-binding and natural transformation in V. cholerae

As mentioned above, prior work has established that two positively charged residues at the C-terminus of the minor pilin FimT contribute to competence T4P DNA-binding and natural transformation in a number of diderms [2, 19–21]. Specifically, work in *V. cholerae* highlighted one FimT residue (FimT_Vc_ ^R154^) that was critical for DNA-binding and natural transformation [2]. Subsequent studies in *Legionella pneumophila*, *Xyllela fastidiosa*, and *Acinetobacter baylyi* highlighted an additional C-terminal residue [19–21] and a broadly conserved G-[R/K]-X-[R/K] motif at the C-terminus of FimT where the two R/K residues play a critical role in DNA-binding (**Fig. S1**).

To recapitulate these previous findings, we assessed the impact of the R/K residues in the FimT_Vc_ G-[R/K]-X-[R/K] motif (FimT_Vc_ ^R154^ and FimT_Vc_ ^K156^) on natural transformation and competence T4P-dependent DNA-binding in *V. cholerae*. We mutated these residues to glutamine, which has a neutral charge while maintaining a similar size to the native R/K residues, because we aimed to abolish the charge of these residues while minimizing the impact of the mutation on overall protein folding and structure. We found that *fimT_Vc_ ^R154Q^* reduced NT to the limit of detection, similar to Δ*fimT*_Vc_. While *fimT_Vc_ ^K156Q^* transformed better than Δ*fimT*_Vc_, this mutant still exhibited a 4-log decrease in NT compared to the parent (**Fig. 1A**). Thus, these residues play a major role in NT.

**Fig. 1.**
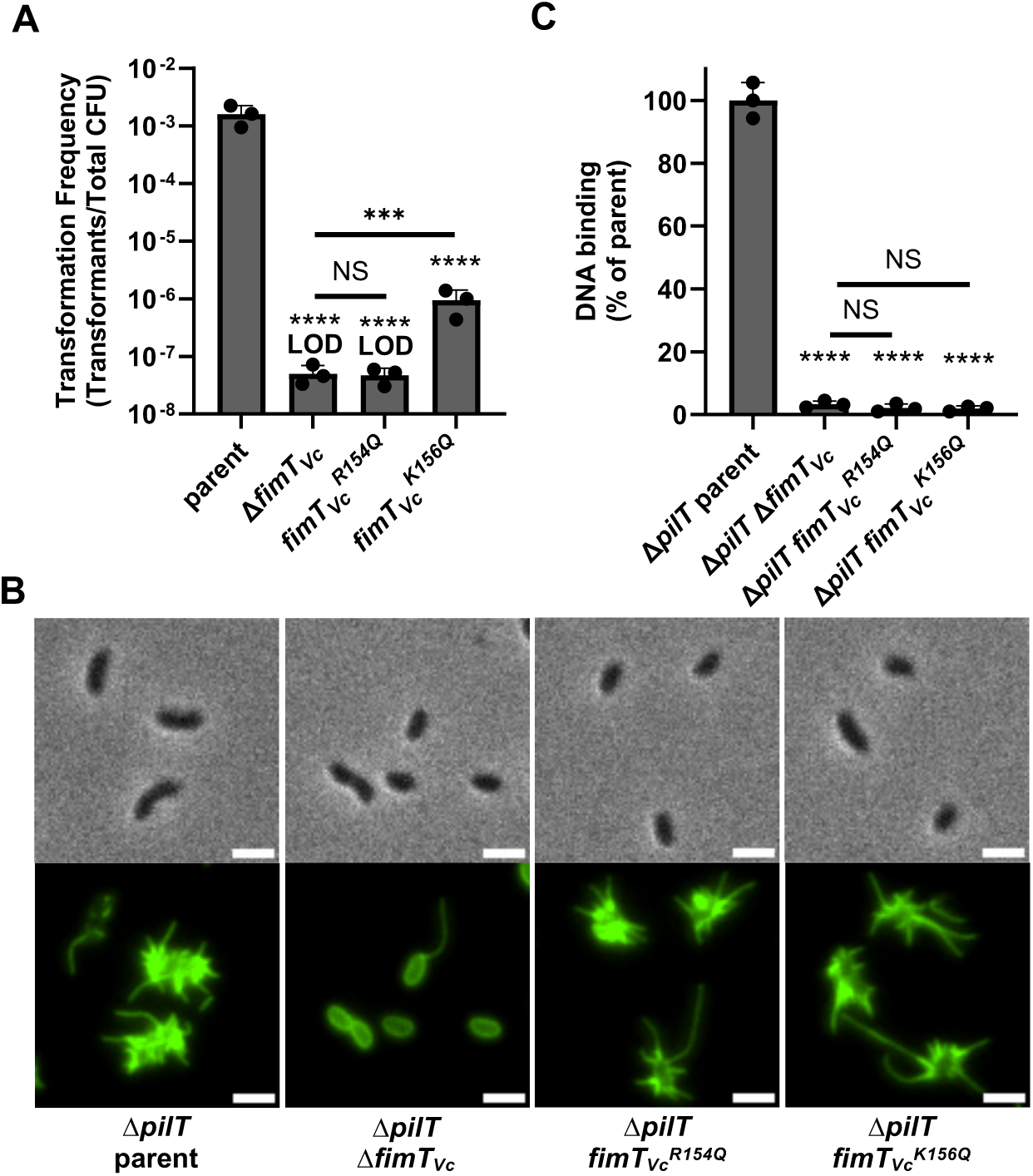
The FimT*_Vc_* G-[R/K]-X-[R/K] motif is critical for natural transformation and DNA-binding in *V. cholerae*. (**A**) Natural transformation assay of the indicated *V. cholerae* strains. (**B**) Representative images of labeled competence T4P in the indicated *V. cholerae* strains. Scale bars, 2 µm. (**C**) DNA-binding assay of the indicated *V. cholerae* strains. Data in **A** and **C** are from at least 3 independent biological replicates and shown as the mean ± SD. Images in **B** are representative of at least 3 independent biological replicates. Statistical comparisons were made by one-way ANOVA with Turkey’s multiple comparison test of the log-transformed data. NS, no significance; *** = *p* < 0.001; **** = *p* < 0.0001. LOD, limit of detection. Statistical identifiers directly above bars represent comparisons to the parent.

The minor pilin FimT_Vc_ is critical for pilus assembly in *V. cholerae* [25], likely because it is a member of the tip-associated complex that initiates pilus assembly. Thus, a trivial explanation for the observed reduction in NT is that these mutations in *fimT_Vc_* disrupt competence T4P assembly. To test this, we used a previously described approach to directly label competence T4P using a strain background where the major pilin contains a cysteine mutation (*pilA^S67C^*) that allows for labeling pili with a fluorescently-conjugated maleimide dye [26]. Competence T4P in *V. cholerae* are highly dynamic [2, 27], so, to assess piliation defects we deleted the retraction motor ATPase, *pilT*, in these strains. This allows cells to extend pili, but greatly reduces retraction, thus sensitizing our ability to assess competence T4P assembly defects in static images. As expected, the parent strain is hyperpiliated in these experiments, and consistent with FimT_Vc_ playing a critical role in competence T4P assembly, Δ*fimT_Vc_* exhibits an almost complete loss in surface piliation (**Fig. 1B**). By contrast, *fimT_Vc_ ^R154Q^* and *fimT_Vc_ ^K156Q^* are hyperpiliated like the parent (**Fig. 1B**), suggesting that these mutations do not affect competence T4P assembly.

Because *fimT_Vc_ ^R154Q^* and *fimT_Vc_ ^K156Q^* do not exhibit a competence T4P assembly defect, we hypothesize that the reduction in NT in these mutants is due to a defect in competence T4P DNA-binding activity. To test this directly, we assessed the ability of hyperpiliated cells (i.e., Δ*pilT* strains) to pulldown fluorescently labeled DNA using a previously established assay [2]. When we perform this experiment, we find that *fimT_Vc_ ^R154Q^* and *fimT_Vc_ ^K156Q^* exhibit a reduction in DNA-binding that is indistinguishable from Δ*fimT_Vc_* (**Fig. 1C**). Together, these results strongly suggest that the R/K residues in the FimT_Vc_ G-[R/K]-X-[R/K] motif directly contribute to competence T4P-dependent DNA-binding in the diderm *V. cholerae*.

### Structural modeling reveals homology between the minor pilins that compose the competence T4P tip complexes in monoderms and diderms

Next, we sought to determine if the molecular basis for DNA-binding by monoderm competence T4P was similar to what is observed in diderms. Simple BLAST searches failed to reveal a FimT_Vc_ homolog in the monoderm *S. pneumoniae*. Work that was published while this study was underway, however, defined the minor pilins from *Streptococcus sanguinis* and demonstrated that these proteins had strong predicted structural similarity to the minor pilin tip complexes of diderm T4P and T2SSs [6].

Because of the predicted structural similarity between the tip complexes in monoderm and diderm T4P, we sought to use structural predictions to define the minor pilin homologs between *V. cholerae* and *S. pneumoniae*. Specifically, we used AlphaFold-multimer (AF-m) to generate high-confidence models of the minor pilin tip complexes (**Fig. S2A-B**). Consistent with prior work [6], we found a high degree of predicted structural homology between these T4P tip complexes despite a lack of sequence homology.

There are several factors that suggest that these AF-m models are physiologically relevant. First, the minor pilins are arranged in a right-handed helix, which is consistent with the solved structures of T4P filaments in monoderms and diderms [28–31]. Second, a genetic signature unique to the minor pilin that caps the tip complex is present. All pilins other than the capping pilin require a conserved E5 residue (position in mature pilin) [32], which neutralizes the charge of the N-terminus of the previously added pilin [31]. Because the capping pilin is the first pilin added, it does not require this E5 residue. Both *V. cholerae* and *S. pneumoniae* have one minor pilin that naturally lacks E5, and in both AF-m models, this pilin is predicted to be the capping pilin. Third, the arrangement of the minor pilins in these AF-m models reflects the biochemical necessity for the E5 residue to neutralize the N-terminus of the previous pilin in the complex (**Fig. S2C**).

Because AF-m predicted that the competence T4P tip complexes of diderms and monoderms were similar, we hypothesized that each minor pilin in *V. cholerae* had a structural homolog in *S. pneumoniae*. As mentioned above, the absence of the E5 residues defined the capping pilin as PilX_Vc_ and ComGG in *V. cholerae* and *S. pneumoniae*, respectively. In *V. cholerae*, the helical assembly of the minor pilins within the tip complex starting at the capping pilin is PilX_Vc_ ➔ PilV_Vc_ ➔ PilW_Vc_ ➔ FimT_Vc_ (**Fig. S2B**). In *S. pneumoniae*, the order of minor pilins from the capping pilin is ComGG ➔ ComGE ➔ ComGF ➔ ComGD (**Fig. S2B**). If the structural homologs of each minor pilin are similarly arranged in the *S. pneumoniae* tip complex model, this would suggest that ComGD = FimT_Vc_, ComGF = PilW_Vc_, and ComGE = PilV_Vc_. To test this, we aligned the headgroups of each *S. pneumoniae* minor pilin to the headgroups of each *V. cholerae* minor pilin (**Fig. S3**). Consistent with our hypothesis, ComGD most closely aligns with FimT_Vc_, ComGF most closely aligns with PilW_Vc_, and ComGE most closely aligns with PilV_Vc_ (**Fig. S3**). This structural arrangement of minor pilins within the tip complex is likely also conserved in the T2SS [24, 33, 34], a member of the broader type IV filament family, which further supports these minor pilin designations (**Fig. S4)**. The minor pilins of the *S. pneumoniae* competence T4P, *V. cholerae* competence T4P, and *Pseudomonas aeruginosa* T2SS are all arranged in a single operon (**Fig. S5**). Interestingly, while the structural arrangement of the minor pilin proteins is conserved across these systems, the genetic arrangement of the minor pilin genes is not (**Fig. S4, S5**); with the *V. cholerae* competence T4P minor pilins exhibiting a distinct arrangement. While it is not clear what led to this difference, this distinction suggests these systems have diverged evolutionarily. Regardless, to reflect the observed conservation in structure between the minor pilins of these systems, and to ease comparisons, moving forward we will refer to *S. pneumoniae* minor pilins as follows: ComGG = PilX_Spn_, ComGE = PilV_Spn_, ComGF = PilW_Spn_, and ComGD = FimT_Spn_. *The G-[R/K]-X-[R/K] motif is conserved* in *FimT_Spn_ but is not critical for natural transformation in S. pneumoniae* Our results in *V. cholerae* indicated that FimT*_Vc_* ^R154^ and FimT*_Vc_* ^K156^ are critical for DNA-binding and NT. When we align the FimT*_Vc_* and FimT*_Spn_* AF-m models, these positively charged residues are predicted to be conserved in FimT_Spn_ (FimT_Spn_^K125^ and FimT_Spn_^K127^; **Fig. 2A**), forming the same G-[R/K]-X-[R/K] DNA-binding motif present in many diderm FimT homologs. Interestingly, this similarity in predicted structure is not revealed by multiple sequence alignments (MSAs), likely due to poor conservation at the sequence level (**Fig. S6**).

**Fig. 2.**
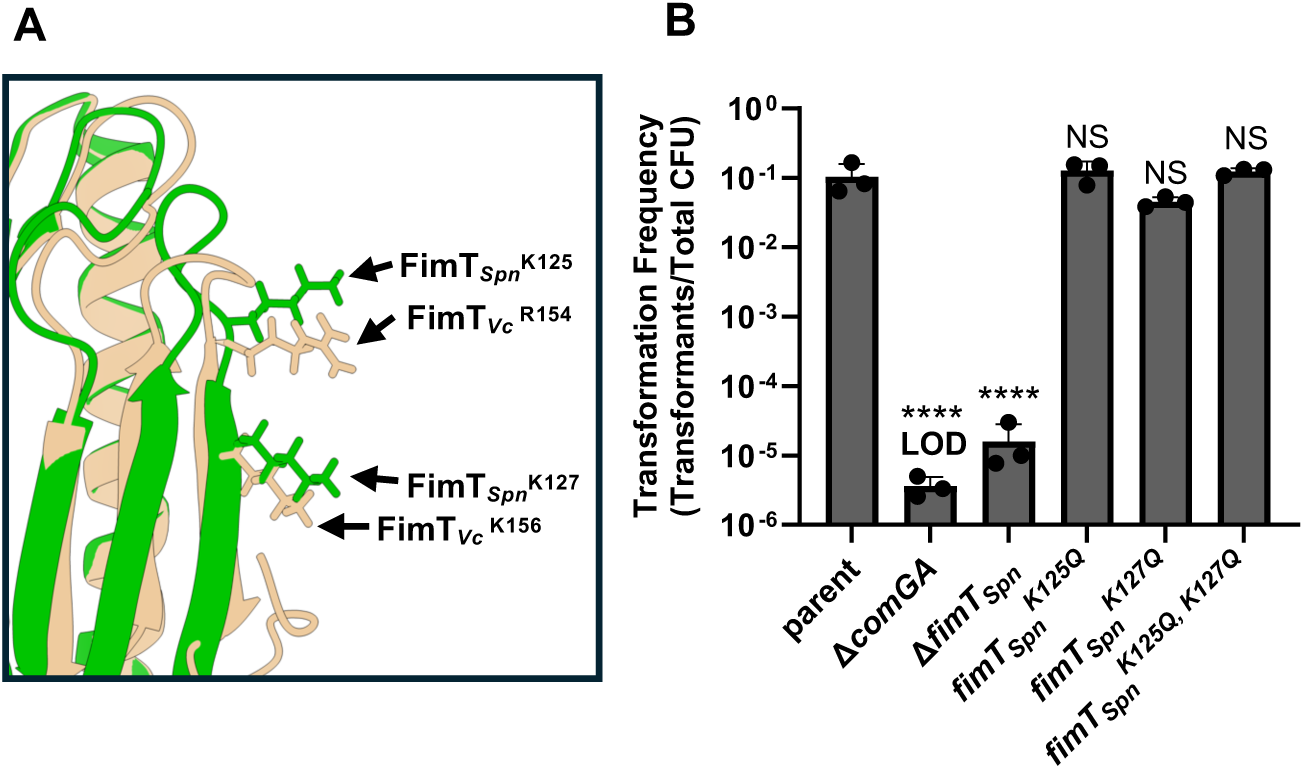
Positively charged residues in the FimT*_Spn_* G-[R/K]-X-[R/K] motif are not critical for NT in *S. pneumoniae*. (**A**) AF-m models of FimT*_Vc_* (tan) and FimT*_Spn_* (green) were aligned and the conserved R/K residues in the G-[R/K]-X-[R/K] motif are highlighted. (**B**) Natural transformation assay of the indicated *S. pneumoniae* strains. Data in **B** are from at least 3 independent biological replicates and shown as the mean ± SD. Statistical comparisons were made by one-way ANOVA with Turkey’s multiple comparison test of the log-transformed data. NS, no significance; **** = *p* < 0.0001. LOD, limit of detection. Statistical identifiers directly above bars represent comparisons to the parent.

Thus, we hypothesized that the molecular basis of DNA-binding may be broadly conserved between monoderm and diderm competence T4P.

To test this, we assessed the impact of mutating these residues in FimT*_Spn_* on natural transformation in *S. pneumoniae*. While the parent is highly transformable in NT assays, a mutant that cannot assemble pili (Δ*comGA*; encoding the competence T4P motor ATPase) and Δ*fimT_Spn_* exhibit an ∼4-log decrease in NT (**Fig. 2B**). By contrast, there was no significant reduction in NT in either *fimT_Spn_^K125Q^* or *fimT_Spn_^K127Q^* compared to the parent (**Fig. 2B**). We hypothesized that these residues may be redundant for DNA-binding. However, even when both residues are mutated (*fimT_Spn_^K125Q, K127Q^*), NT is indistinguishable from the parent (**Fig. 2B**). This suggests that while the G-[R/K]-X-[R/K] motif is conserved between FimT_Vc_ and FimT_Spn_, its function in DNA-binding may not be conserved.

*Positively charged residues in FimT_Spn_ and PilW_Spn_ facilitate natural transformation in S. pneumoniae* Because the positively charged residues in the FimT_Spn_ G-[R/K]-X-[R/K] motif did not impact NT, we sought to identify minor pilin residues that contribute to NT to uncover the molecular basis of DNA-binding in monoderm competence T4P. As mentioned previously, competence T4P-dependent DNA-binding likely relies on electrostatic interactions. So, we took an unbiased approach to identify positively charged residues in the *S. pneumoniae* minor pilin tip complex that might contribute to DNA-binding by analyzing the electrostatic surface map of the AF-m model.

From this analysis, we first noticed that the extended C-terminal tail of the capping minor pilin, PilX_Spn_, contains a large number of lysine residues. To test if these positively charged residues contribute to NT, we generated mutants of PilX_Spn_ to either (1) delete this extended C-terminal region or (2) to mutate all lysine residues to glutamine. NT of these mutants, however, were indistinguishable from the parent (**Fig. S7**), which suggests that the C-terminal domain of PilX_Spn_ is not critical for DNA-binding.

Aside from the charged C-terminus of PilX_Spn_, we also identified a large patch of positively charged residues in the tip complex that spanned two minor pilins – FimT_Spn_ (FimT_Spn_^K105^, FimT_Spn_^R116^, FimT_Spn_^K125Q^, and FimT_Spn_^K127Q^) and PilW_Spn_ (PilW_Spn_^R102^, PilW_Spn_^K103^, PilW_Spn_^R107^, and PilW_Spn_^R109^) – that includes the G-[R/K]-X-[R/K] motif mentioned above (**Fig. 3A**). This is in stark contrast to the electrostatic surface map of the *V. cholerae* competence T4P tip complex, where strong positive charge is only observed around the G-[R/K]-X-[R/K] motif in FimT*_Vc_* (**Fig. 3A**).

**Fig. 3.**
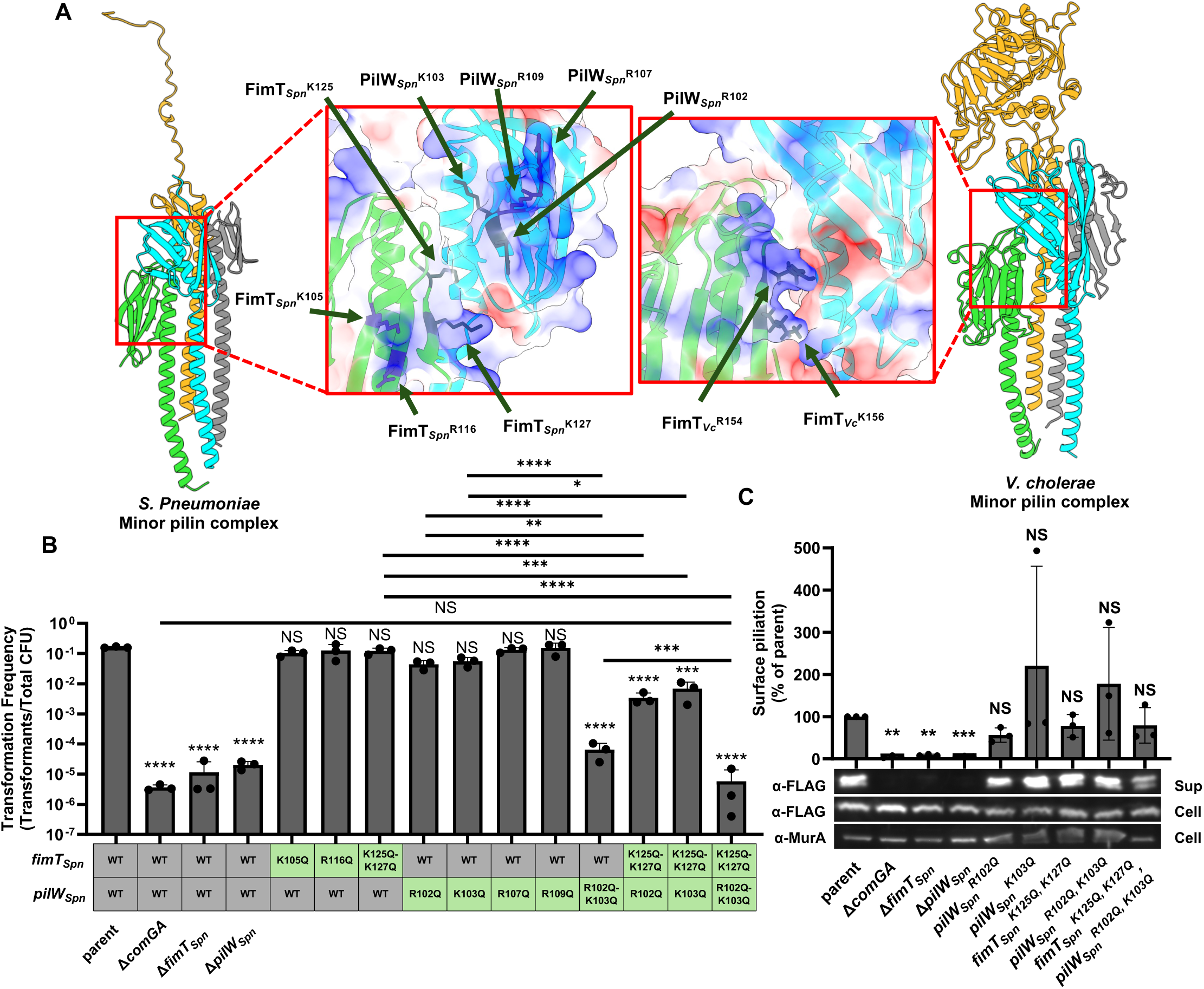
Model guided mutagenesis reveals positively charged residues in FimT*_Spn_* and PilW*_Spn_* that are critical for NT in *S. pneumoniae*. (**A**) AF-m models of the indicated minor pilin tip complexes. Insets highlight the electrostatic surface map of each complex and the positively charged residues targeted for mutagenesis. (**B**) NT assay of the indicated *S. pneumoniae* strains. (**C**) Representative western blot and quantification of ComGC-FLAG (α-FLAG; to assess ComGC levels) and MurA (α-MurA; loading control) in whole cell lysates and sheared supernatants from the indicated ComGC-FLAG *S. pneumoniae* strains. Surface piliation is defined as the ComGC-FLAG density in the sup normalized to the MurA loading control. Results from each replicate were then normalized to the parent and plotted. Data in **B** and **C** are from at least 3 independent biological replicates and shown as the mean ± SD. Blots in **C** are representative of 3 independent biological replicates. Statistical comparisons in **B** were made by one-way ANOVA with Turkey’s multiple comparison test of the log-transformed data and comparisons in **C** were made by one-sample Student’s t-test of the log-transformed data. NS, no significance; * = *p* < 0.05; ** = *p* < 0.01; *** = *p* < 0.001; **** = *p* < 0.0001. Statistical identifiers directly above bars represent comparisons to the parent.

We hypothesized that the residues that make up this positive patch contribute to DNA-binding and uptake during NT. To test this, we mutated each residue to glutamine to assess their role during NT. Mutation of no individual residue had a significant impact on NT, however, *pilW_Spn_^R102Q^* and *pilW_Spn_^K103Q^* exhibited a slight, but non-significant decrease in NT (**Fig. 3B**). When these two residues in PilW_Spn_ were mutated together (*pilW_Spn_^R102Q, K103Q^*), an ∼3-log decrease in NT was observed (**Fig. 3B**), suggesting that these residues were synergistically epistatic (*i.e.*, the phenotype of the double mutant is greater than the combined deficits of each single mutant). This suggests that these residues are redundant in their ability to promote NT. These results suggest that these positively charged residues in PilW_Spn_ may play a major role in DNA-binding in monoderm competence T4P. Interestingly, the positively charged residues we found in PilW_Spn_ were not conserved in the PilW homologs of diderms (**Fig. S8**).

The PilW_Spn_^R102^ and PilW_Spn_^K103^ residues are predicted to be in close proximity to the FimT_Spn_^K125^ and FimT_Spn_^K127^ residues within the tip complex (**Fig. 3A**). Thus, we hypothesized that FimT_Spn_ and PilW_Spn_ may work together to facilitate DNA-binding. To test this, we combined the *fimT_Spn_^K125Q,K127Q^* mutation, which exhibited no loss of NT, with either *pilW_Spn_^R102Q^* or *pilW_Spn_^K103Q^*. We found that both the *fimT_Spn_^K125Q,K127Q^ pilW_Spn_^R102Q^* and *fimT_Spn_^K125Q,K127Q^ pilW_Spn_^K103Q^* mutants exhibited an ∼2-log decrease in NT (**Fig. 3B**), indicating that these mutations are synergistically epistatic, which is consistent with these residues working together to facilitate NT. Consistent with cooperative DNA-binding by FimT_Spn_ and PilW_Spn_, we found that NT of *fimT_Spn_^K125Q, K127Q^ pilW_Spn_^R102Q, K103Q^* was significantly reduced compared to *fimT_Spn_^K125Q, K127Q^* or *pilW_Spn_ ^R102Q, K103Q^*. Furthermore, the phenotype of *fimT_Spn_^K125Q, K127Q^ pilW_Spn_^R102Q, K103Q^* was indistinguishable from Δ*comGA* (a T4P assembly mutant), indicating that the combined activity of these residues in FimT_Spn_ and PiW_Spn_ are critical for NT.

In addition to their potential role in DNA-binding, these minor pilins are also critical for T4P assembly [6]. Thus, a trivial explanation for the NT phenotypes observed is that the residues mutated in FimT_Spn_ and PilW_Spn_ simply diminish T4P assembly. To test this, we used mechanical shearing to detect the assembly of surface pili by western blot analysis for the competence T4P major pilin, ComGC, as previously described [35]. To facilitate detection of ComGC in these assays, all strains harbored a previously described functional ectopic FLAG-tagged allele of ComGC (ComGC-FLAG) [35]. ComGC-FLAG was detected in sheared supernatants to assess surface assembly and in the total cell lysates to ensure expression. Cell lysates were also probed for MurA as a loading control. In this assay, the parent strain exhibits strong ComGC-FLAG signal in sheared supernatants, which is consistent with surface assembly of T4P in this background (**Fig. 3C**). An assembly deficient mutant, Δ*comGA*, showed almost no ComGC-FLAG in the sheared supernatant but abundant signal in the cell fraction, which is consistent with the lack of pilus assembly in this mutant (**Fig. 3C**). Similarly, surface piliation was undetectable in Δ*fimT_Spn_* and Δ*pilW_Spn_*, which is consistent with their established role in pilus assembly [6]. By contrast, surface piliation was indistinguishable from the parent for all of the *fimT_Spn_* and *pilW_Spn_* point mutants tested, including *fimT_Spn_^K125Q, K127Q^ pilW_Spn_^R102Q, K103Q^* (**Fig. 3C**). Because these mutations do not affect T4P biogenesis, this suggests that the NT deficits observed are likely due to a decrease in DNA-binding activity.

### Positively charged residues in FimT_Spn_ and PilW_Spn_ are critical for competence T4P DNA-binding in S. pneumoniae

Thus far, our results demonstrate that four positively charged residues (FimT_Spn_^K125,K127^ and PilW_spn_^R102,K103^) in the minor pilin tip complex are critical for NT. Because these mutations do not affect competence T4P biogenesis, we hypothesized that they are required for DNA-binding. Unfortunately, the whole cell DNA-pulldown assay used to assess *V. cholerae* competence T4P-dependent DNA-binding cannot be used to test this question because competence T4P in *S. pneumoniae* are highly dynamic and the molecular mechanism underlying T4P retraction remains unclear. Thus, we developed an assay to directly assess the DNA-binding activity of sheared *S. pneumoniae* competence T4P.

Specifically, ComGC-FLAG pili are first mechanically sheared from the surface of competent *S. pneumoniae* cells and then captured on anti-FLAG magnetic beads. The captured pili are then incubated with DNA to allow for binding to occur and then washed to remove unbound DNA. Beads are then eluted with FLAG peptide to release the bound T4P and DNA. The eluate is then evaluated (1) by western blotting for ComGC-FLAG to assess the capture of pili and (2) by quantitative PCR to determine the amount of bound DNA (see **Methods** for details).

When we performed this assay, we found that competence T4P from the parent robustly bound DNA. This is in stark contrast to the poor DNA-binding by Δ*comGA* (**Fig. 4A**). This is consistent with the inability of Δ*comGA* to assemble competence T4P, as supported by the absence of ComGC-FLAG in pulldown fractions when compared to the parent (**Fig. 4B**). This result also demonstrates that the ComGC-FLAG signal in pulldown fractions can be attributed to assembled competence T4P. When we assessed *fimT_spn_^K125Q, K127Q^ pilW_Spn_ ^R102Q,K103Q^* in this assay, we found that while the amount of competence T4P pulled down was similar to the parent (**Fig. 4B**), DNA-binding was reduced to the background level observed in Δ*comGA* (**Fig. 4A**). These results demonstrate that the molecular basis for DNA-binding by *S. pneumoniae* competence T4P relies on these positively charged residues in FimT_Spn_ and PilW_Spn_.

**Fig. 4.**
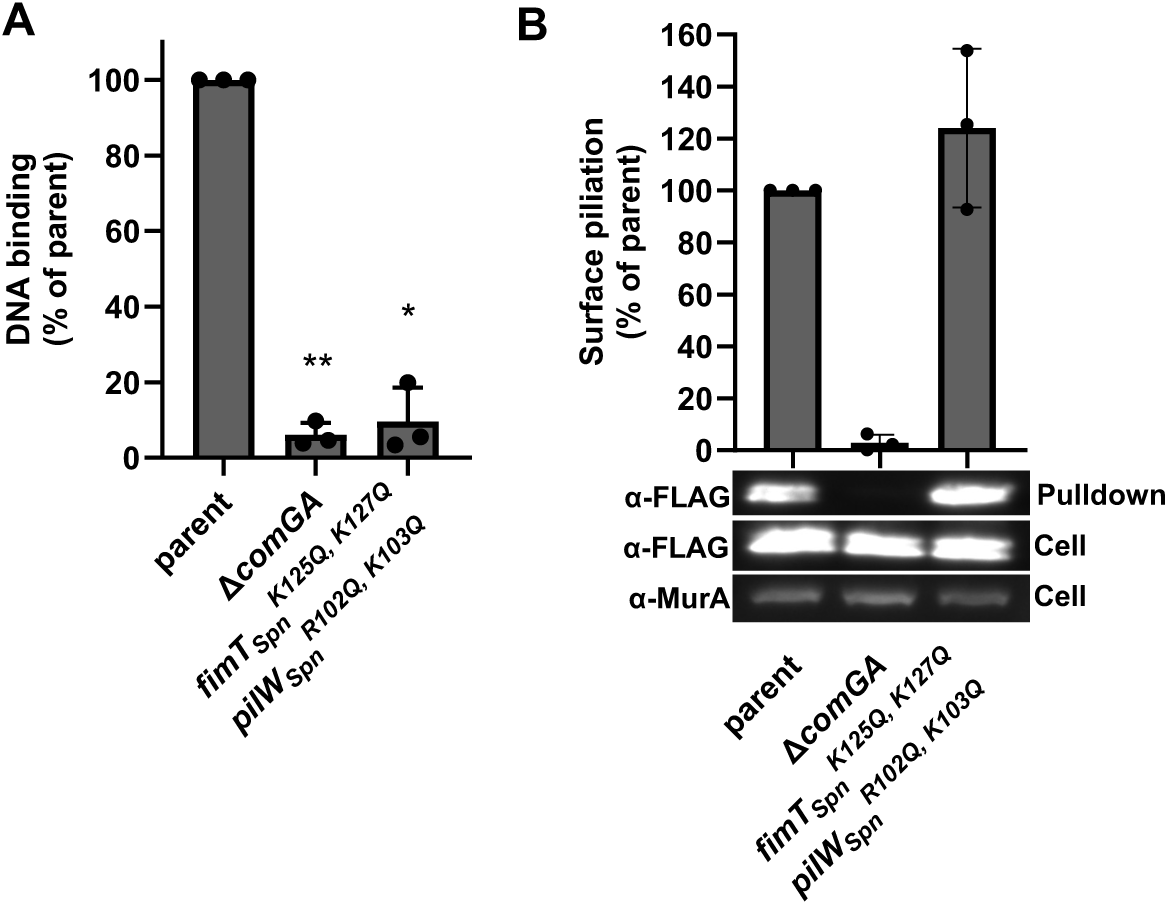
The *fimT^K125Q,K127Q^ pilW^R102Q,K103Q^* mutation eliminates the DNA-binding activity of *S. pneumoniae* competence T4P. (**A**) DNA binding assay using competence T4P captured from sheared supernatants of the indicated *S. pneumoniae* strains. (**B**) Representative western blot and quantification of ComGC-FLAG (α-FLAG; to assess ComGC expression) and MurA (α-MurA; loading control) in whole cell lysates and purified T4P from sheared supernatants (pulldown) as indicated. Data in **A** and **B** are from at least 3 independent biological replicates and shown as the mean ± SD. Statistical comparisons were made by one-sample t test of the log-transformed data. * = *p* < 0.05; ** = *p* < 0.01. Statistical identifiers directly above the bar represent comparisons to the parent.

### Positively charged residues in PilW_Spn_ are conserved in monoderms

Our results suggest that the molecular basis for DNA binding by *S. pneumoniae* competence T4P is distinct from what has been described in diderm organisms [2, 19–21]. Specifically, we show that in *S. pneumoniae*, positively charged residues in PilW_Spn_ play the dominant role in DNA-binding, while in diderms, this activity is primarily driven by positively charged residues in FimT.

Next, we sought to determine whether this was a unique property of *S. pneumoniae*, or if PilW-dependent DNA-binding is a conserved property of monoderm competence T4P. To test this, we first aligned the amino acid sequences from *S. pneumoniae*, *Bacillus subtilis*, *Streptococcus sanguinis*, and *Lactococcus lactis* (**Fig. S9A**). Three of four organisms maintained the complete G-[R/K]-X-[R/K] motif in FimT (*B. subtilis* partially maintained this motif) and all four organisms maintained the RK residues in PilW. We then used AlphaFold3 (AF3) to model the competence T4P minor pilin tip complexes of all four organisms (**Fig. S9B**). Each complex maintained (1) the right-handed helical structure, (2) the protein arrangement, and (3) the gene locus architecture observed in *S. pneumoniae*.

The *S. pneumoniae, S. sanguinis,* and *L. lactis* competence minor pilin complex models had nearly identical structures, with all four DNA-binding residues conserved (**Fig. S9**). The two positively-charged residues in PilW_Spn_ required for DNA-binding (PilW_Spn_^R102,K103^) are also conserved in the *B. subtilis* PilW homolog (PilW_Bs_ ^R70,K71^), while only one of the two positively-charged FimT_Spn_ residues is conserved (**Fig. S9**). The stronger conservation of positively charged residues in PilW compared to FimT homologs among monoderms supports the hypothesis that PilW plays the dominant role in DNA-binding in monoderm competence T4P.

## DISCUSSION

This study sheds light on the molecular basis of DNA binding by monoderm competence T4P. Specifically, we show that four positively charged residues that span two minor pilins, FimT and PilW, contribute to NT and DNA-binding in *S. pneumoniae*. Furthermore, we show that these residues are conserved in other monoderms, but not diderms. These results are notable for two reasons: (1) they suggest that the molecular basis for DNA-binding is distinct in monoderm and diderm competence T4P, and (2) they suggest that DNA-binding by competence T4P is a property of the tip-associated complex of minor pilins, not a feature of a single minor pilin. We discuss each of these points in detail below.

The FimT G-[R/K]-X-[G/K] motif has been shown to play a critical role in DNA-binding among diderm competence T4P [2, 19, 20]. While this motif is conserved in many monoderm FimT homologs, we show that it, surprisingly, plays a relatively minor role during NT in *S. pneumoniae*. Instead, we found that the neighboring minor pilin, PilW, has two positively charged residues that are important for NT in *S. pneumoniae*. Furthermore, we show that these PilW residues are conserved among monoderm competence T4P and are not conserved in the PilW homologs of diderms. These data suggest that competence T4P in monoderms may rely more heavily on PilW for DNA-binding during NT, while diderms rely more heavily on FimT. The high degree of conservation among T4P systems [36] suggests that competence T4P likely shared a common ancestor. Thus, our results suggest that the molecular basis for DNA-binding has diverged between monoderm and diderm competence T4P. These differences could simply represent genetic drift. Or it is tempting to speculate that this divergence reflects distinct evolutionary pressures. T4P-dependent DNA uptake in diderms only requires pilus-bound DNA to traverse through an outer membrane secretin pore to enter the periplasm; while in monoderms, the T4P must weave bound DNA through the thick cell wall. Thus, constraints on the DNA-binding activity of competence T4P may differ among monoderms and diderms to accommodate these distinct requirements.

A number of prior studies have aimed to define the molecular basis of competence T4P DNA-binding by biochemically characterizing individual pilins. While these studies have been highly valuable at defining important features of these pilins, they have led to the notion that the DNA-binding observed by individual pilins is reflective of the DNA-binding mechanism of intact competence T4P. This is perhaps best supported by data in *Neisseria*, where DNA is bound in a sequence-specific manner by the minor pilin ComP. Biochemical analysis of the purified headgroup of ComP demonstrated that this minor pilin is sufficient to bind to *Neisseria* DNA-uptake sequences (DUSs) *in vitro*. And detailed structure-function analysis revealed that mutations that diminish DNA-binding of ComP *in vitro*, directly correlated with a reduction in NT *in vivo* (*i.e.*, a 1-log reduction in DNA-binding *in vitro* yields a 1-log deficit in NT *in vivo*) [14, 37, 38]. This evidence strongly supports the model that ComP is necessary and sufficient for the uptake of sequence-specific DNA (i.e., DNA containing a DUS) in *Neisseria* species. ComP-dependent DNA uptake by *Neisseria* competence T4P, however, is an outlier among naturally transformable microbes.

The uptake of DNA in almost every other competent species (aside from *Haemophilus spp.*) occurs in a sequence-independent manner. There are a number of observations among these sequence-independent systems that contradict the model that competence T4P DNA-binding can be elucidated through the biochemical analysis of individual minor pilins. Biochemical characterization of major and minor pilins from *Thermus thermophilus* (ComZ, PilA1, PilA2, PilA3) [39, 40], *Clostridioides difficile* (PilJ, PilW) [41], and *X. fastidiosa* (FimT3) [20] has demonstrated that a number of architecturally distinct pilins bind to DNA *in vitro*. While some of these studies identified residues that diminish DNA-binding *in vitro*, the impact of these residues on DNA-binding and/or NT *in vivo* was not tested. So, the physiological relevance of the DNA-binding observed for these purified pilins *in vitro* remains unclear. Also, in *L. pneumophila*, while mutation of the G-[R/K]-X-[G/K] motif reduced DNA-binding by purified FimT_Lp_ ∼1-log *in vitro*, mutating these residues *in vivo* decreased NT >3- logs to the limit of detection [19]. This discrepancy suggests that the DNA-binding observed by purified FimT *in vitro* may not accurately reflect the DNA-binding of intact competence T4P. Thus, for sequence-independent competent species, it remains unclear whether the biochemical properties of individual minor pilins can recapitulate the DNA-binding activity of intact competence T4P.

In this study, we provide strong genetic evidence that positively charged residues in both FimT_Spn_ and PilW_Spn_ work together to facilitate DNA-binding and NT in *S. pneumoniae*. This is also supported by prior work in *V. cholerae*, where it was shown that one positively charged residue in PilW_Vc_ (PilW_Vc_ ^K144Q^) plays a minor role in DNA-binding [2]. This highlights that the DNA-binding activity of competence T4P is likely not dictated by a single minor pilin but is instead a property of the minor pilin tip complex. Thus, studies that seek to understand the biochemical properties of DNA-binding by competence T4P should either focus on (1) analyzing intact T4P filaments or (2) by reconstituting the minor pilin tip complex *in vitro*. Reconstitution of a minor pilin tip complex has not been accomplished in any T4P system to date. This is because the hydrophobic tails of intact pilins, which likely contribute to the formation and stability of the tip complex, make them incredibly difficult to purify and characterize *in vitro*. Also, it is possible that minor pilin interactions with the T4P machine are required for proper assembly, further complicating reconstitution of the minor pilin tip complex *in vitro*.

Altogether, our results help elucidate how monoderm competence T4P bind DNA to facilitate a widely conserved mechanism of horizontal gene transfer. T4P are nearly ubiquitous surface appendages that facilitate diverse activities in bacterial species including twitching motility, adherence, virulence, and protein secretion.

Recent work highlights that these activities also rely on minor pilins [11, 12, 42–45]. As mentioned above, minor pilin tip complexes have not been biochemically reconstituted in any system. Our results suggest that these tip complexes exhibit properties that cannot be recapitulated through the characterization of individual minor pilins. Thus, this study also highlights how structural modeling can be successfully employed to dissect the molecular mechanisms of these biochemically intractable complexes *in situ*.

## METHODS

### Bacterial strains and culture conditions

All *V. cholerae* strains were derived from *V. cholerae* E7946 [46]. *V. cholerae* cells were routinely grown in LB Miller broth rolling at 30⁰C or on LB miller agar at 30⁰C supplemented with trimethoprim (10 µg/mL), spectinomycin (100 µg/mL), kanamycin (50 µg/mL), or zeocin (100 µg/mL) as appropriate. All *S. pneumoniae* strains were derived from RL001, an R6 strain from the Fronzes lab [35]. *S. pneumoniae* cells were routinely grown in Casamino Acid Tryptone medium with 0.2% glucose and 16 mM K_2_HPO_4_ (CAT+GP) statically at 37⁰C or on Sheep’s blood agar (ThermoFisher) at 37⁰C in 5% CO_2_ (candle jar extinction method) supplemented with spectinomycin (100 µg/mL) or erythromycin (0.3 µg/mL) as appropriate.

### Construction of mutant strains

All strains were generated by natural transformation using splicing-by-overlap-extension (SOE) PCR mutant constructs exactly as previously described [47]. All strains were confirmed by PCR and/or sequencing. For a detailed list of mutants in this study, see **Table S1**. For a complete list of primers used to generate mutant constructs, see **Table S2**.

### Natural transformation assays

All *V. cholerae* and *S. pneumoniae* strains used in this study were engineered to bypass the native signals required for competence development.

To bypass native competence development in *V. cholerae*, the master competence regulator, TfoX, was overexpressed via an isopropyl β-d-1-thiogalactopyranoside (IPTG)-inducible P*_tac_* promoter and cells were locked in a high cell density state via deletion of *luxO*, as previously described [47]. Chitin-independent transformation assays to determine transformation frequency were performed as previously described [2]. Briefly, cells were grown at 30⁰C rolling in LB supplemented with 100 µM IPTG, 20 mM MgCl_2_, 10 mM CaCl_2_ to late log. Then ∼10^8^ CFUs were diluted into 350 µL of instant ocean medium (7g/L; Aquarium Systems) and incubated on a 30⁰C dry heat block (USA Scientific). Next, ∼200 ng of transforming DNA (tDNA) with a trimethoprim resistance cassette (a ΔVC0462::Tm^R^ PCR product) was added into each reaction. After tDNA addition, reactions were incubated statically at 30⁰C for exactly 5 minutes, and then 2 µL of DNase I (2000 U/mL, NEB) was added to degrade any remaining exogenous tDNA. To allow for tDNA integration into the genome, reactions were incubated statically at 30⁰C overnight. Reactions were outgrown by adding 500 µL of LB and shaking at 37⁰C for 2 hours, and then plated for quantitative culture on LB agar supplemented with trimethoprim (transformants), and plain LB agar (total viable counts). Transformation frequency is defined as CFU/mL of transformants divided by the CFU/mL of total viable counts. Additionally, control reactions were performed for each strain tested where no tDNA was added.

To bypass native competence in *S. pneumoniae*, strains contained a deletion of *comC*, which encodes the competence stimulating peptide (CSP) [35]. Thus, this strain relies on exogenous CSP for competence induction. Strains were first struck out onto Sheep’s blood agar from a frozen stock and grown overnight at 37⁰C in 5% CO_2_. Cells were lifted from blood plates with CAT and diluted to OD_600_ of 0.005 in CAT+GP medium and incubated statically at 37⁰C. Once the culture reached an OD_600_ of ∼0.05, an aliquot of culture was placed in a fresh eppi and supplemented with 1 mM CaCl_2_, 0.4% BSA, and 100 ng/mL of CSP (ThermoFisher).

Reactions were incubated statically on a 37⁰C dry heat block (USA Scientific) for 12 minutes to induce competence. Next, ∼200 ng of tDNA with a spectinomycin resistance cassette (a Δspr0857::Spec^R^ PCR product) was added to each reaction. After tDNA addition, reactions were incubated statically at 37⁰C for exactly 5 minutes, and then 2 µL of DNase I (2000 U/mL, NEB) was added to degrade any remaining exogenous tDNA. Cells were incubated statically at 37⁰C for an additional 55 minutes to allow for tDNA integration and outgrowth. After incubation, reactions were removed from the heat block and plated for quantitative culture on blood agar supplemented with spectinomycin (transformants) and plain blood agar (total viable counts). Transformation frequency is defined as CFU/mL of transformants divided by the CFU/mL of total viable counts. Control reactions were performed for each strain tested where no tDNA was added.

### Microscopy to assess surface piliation of V. cholerae

Cells were grown to induce competence as described above for transformation assays. To observe surface piliation in these assays, all *V. cholerae* strains harbored a Δ*VC0462*::Tm^R^ mutation to inactivate the retraction motor ATPase PilT, which greatly reduces the retraction of extended competence T4P. All *V. cholerae* strains also harbored a *pilA^S67C^* mutation which allows for pilus labeling with a maleimide dye without affecting pilus activity [26]. Once cells had reached late-log, ∼10^8^ cells were harvested and washed in instant ocean medium to remove residual LB. To label competence T4P, washed cells were incubated with 25 ng/mL AlexaFluor 488- maleimide dye (ThermoFisher) for 15 minutes at room temperature. To remove unbound dye, cells were washed three times in instant ocean medium. Then, cells were placed under a 0.4% gelzan pad for imaging. Phase contrast and wide-field fluorescence images were collected using a Nikon Ti-2 microscope using a Plan Apo × 60 objective, a FITC filter cube, a Hamamatsu ORCA Flash 4.0 camera and Nikon NIS Elements imaging software.

### Western blot analysis

*S. pneumoniae* strains were first struck out onto Sheep’s blood agar from a frozen stock and grown overnight at 37⁰C in 5% CO2. Cells were lifted from blood plates with CAT and diluted to an OD_600_ of 0.02 in CAT+GP medium. These cultures were incubated statically at 37⁰C until they reached an OD_600_ of ∼0.2. Cultures were then supplemented with 0.16% BSA, 1mM CaCl_2_, and 100 ng/mL CSP. Cells were incubated at 37⁰C for exactly 7 minutes to allow for competence induction and then placed on ice to halt the progression of competence. Induced cells were centrifuged for 15 minutes at 4⁰C, and then gently washed once in ice-cold “meow mix” (CAT+GP, 1 mM CaCl_2_, 0.2% BSA) to remove dead cell matter, and finally resuspended in 500 µL ice cold “meow mix.” To restart the progression of competence, cells were incubated at 37⁰C (dry heat block) for 6 minutes and then vigorously vortexed for 45 seconds to mechanically shear surface-exposed pili. To promote a second round of T4P extension, cells were incubated at 37⁰C for an additional 5 minutes, and then vigorously vortexed for 45 seconds. To separate cells from the mechanically sheared pili in the supernatants, samples were centrifuged at 4⁰C. To generate samples for whole cell lysates and mechanically sheared T4P, the cell pellet and supernatant fractions were treated as follows.

To prepare whole cell lysates, the cell pellets were resuspended in 500 µL cell lysis buffer [1x FastBreak lysis buffer (Promega), 1% Triton, 1 mM PMSF, 1x protease inhibitor cocktail [0.07 mg/mL phosphoramidon (Santa Cruz), 0.006 mg/mL bestatin (MPbiomedicals/Fisher Scientific), 1.67 mg/mL AEBSF (Gold Bio), 0.07 mg/mL pepstatin A (DOT Scientific), 0.07 mg/mL E64 (Gold Bio; suspended in DMSO)]] and transferred to a flat-capped tube with 0.35 g matrix B (Promega). To complete cell lysis, tubes were beaten on a FastPrep machine (MP biomedicals). Lysates were then centrifuged at 4⁰C and 50 µL of supernatant was mixed 1:1 with 2x SDS-PAGE sample buffer (200 mM Tris pH 6.8, 25% glycerol, 1.8% SDS, 0.02% Bromophenol Blue, 5% β-mercaptoethanol).

Mechanically sheared T4P were concentrated from the supernatant fraction by trichloroacetic acid (TCA) precipitation. Specifically, supernatants were transferred into a new eppi tube containing 5 µL 100 mM PMSF and 5 µL 100x protease inhibitor cocktail to prevent degradation of sheared T4P present in the supernatant. To remove any cells that may have transferred with the supernatant, samples were centrifuged at 4⁰C again. To precipitate sheared pili from the supernatants, supernatant was moved to a new eppi tube containing 30 µL TCA, briefly vortexed, and incubated on ice. Each sample was supplemented with 0.08% BSA during the TCA precipitation to act as an inert carrier. To collect precipitated protein, samples were centrifuged at 4⁰C. The supernatant was completely removed, and the precipitated protein pellet was washed thoroughly in ice-cold acetone. The acetone supernatant was then removed and the protein pellets were placed at 42⁰C to dry.

Protein pellets were then resuspended in 50 µL 0.1 N NaOH and mixed 1:1 with 2x SDS-PAGE sample buffer.

Sheared supernatant samples and whole cell lysate samples were run on 15% SDS-PAGE gels and transferred to PVDF membranes. To detect ComGC-FLAG blots were stained with α-FLAG primary antibodies (Sigma). As a loading control, MurA was detected in whole cell lysates using α-MurA antibodies (courtesy of Malcolm Winkler). Blots were then incubated with HRP-conjugated secondary antibodies, developed using enhanced chemiluminescence (ECL) western blotting substrate (Pierce), and imaged on a ProteinSimple Fluorechem R instrument.

Protein densities were quantified using Fiji imaging software. Supernatant ComGC-FLAG densities were normalized by MurA. Results were then normalized to the parent to combine data from independent blots. The limit of detection was conservatively set to 10% greater than the background intensity.

### DNA-binding assays

To assess DNA binding by *V. cholerae* competence T4P, Δ*pilT* cells were grown to induce competence as described above for transformation assays. For each DNA binding reaction, ∼1×10^8^ cells were harvested, washed twice with binding buffer (1% NaCl, 20mM MgCl_2_, 10mM CaCl_2_, 0.1mg/ml rBSA), and finally resuspended in 50 µL DNA binding buffer. Then, ∼100 ng of Cy3-labeled plasmid was added. DNA was labeled using the Cy3 LabelIt kit (Mirrus Bio) according to manufacturer’s instructions. To allow DNA binding to occur, the reactions were incubated statically at 30⁰C for 15 minutes. Reactions were washed twice with 200 µL of binding buffer and then resuspended in 120 µL of binding buffer. To liberate the Cy3-labeled DNA bound by competence T4P, 2 µL of DNase I was added to each reaction and incubated for 3 minutes at room temperature. To separate cells from the liberated DNA, cells were pelleted by centrifugation. Then, 100 µL of supernatant was collected and placed in a 96-well plate to assess Cy3 fluorescence on a Biotek Synergy H1M plate reader. Data were normalized by the cell density of each reaction. That that end, the cell pellets of each reaction were resuspended in 200 µL of cell buffer (1% NaCl, 20mM MgCl_2_, 10mM CaCl_2_). Then, a 1:10 dilution was made in a 96-well plate and OD_600_ was measured on the plate reader. The Cy3 fluorescence of the supernatant was then divided by the OD_600_ of the corresponding reaction. A “no DNA” control was also prepared to measure background fluorescence associated with *V. cholerae* cells and the buffer. The normalized value for the no DNA control (*i.e.*, background fluorescence) was subtracted from each sample. Results were then normalized to the parent to combine data from independent experiments.

To assess DNA binding by *S. pneumoniae* competence T4P, cells were grown and treated exactly as described above for western blot analysis. Mechanically sheared pili in supernatant fractions were not TCA precipitated, but were instead captured using α-FLAG magnetic beads (Pierce Anti-DYKDDDDK Magnetic Agarose, ThermoFisher). Specifically, supernatants were incubated with nutation for 1 hour at room temperature with α- FLAG beads washed in Tris-buffered saline. Captured T4P were washed twice using a magnetic stand to collect the beads and resuspended in 500 µL CAT+GP with 1 mM PMSF and 1x protease inhibitor cocktail. To allow for T4P DNA-binding, ∼500 ng of heterologous DNA was added and incubated at room temperature with nutation for 1 hour. After this incubation, the beads were washed twice and resuspended in 100 µL of FLAG elution buffer (50 mM Tris pH 7.5, 150 mM NaCl, 1 mM EDTA, 10 mM MgCl2, 0.1% Triton, 2% glycerol) and transferred to PCR tubes. To elute competence T4P (and any associated bound DNA) from beads, 3 µL of 5 mg/mL FLAG peptide (Millipore-Sigma) was added and rotated end over end at room temperature for 20 minutes. Then, 50 µL of the eluate was mixed 1:1 with 2x SDS-PAGE sample buffer and subjected to western blot analysis alongside whole cell lysates as described above.

The remaining 50 µL of eluate was evaluated for captured DNA. The eluate was mixed with 500 µL of PB buffer (5% ethanol, 5.5 M Guanidine HCl, 20 mM Tris pH 6.6) and taken through the steps of PCR purification. DNA was eluted from the column with 30 µL of elution buffer (10 mM Tris pH 8.5) heated to 37⁰C. Eluted fractions were used as template for quantitative PCR (qPCR; Applied Biosystems) to determine the concentration of bound heterologous DNA using the standard curve approach. Results were then normalized to the parent to combine data from independent experiments.

### Structural modeling and model analysis

AlphaFold-multimer was used to generate the majority of the models [48]. We utilized Indiana University’s high-performance computing (HPC) system to run ColabFold [49]. For minor pilin tip complex models, recycles were set to 10 with an early stop tolerance of 0.4. For individual minor pilin models, recycles were set to 30 with an early stop tolerance of 0.2. AlphaFold3 was used to generate models of the *S. pneumoniae*, *S. sanguinis*, *L. lactis*, and *B. subtilis* minor pilin complexes [50] via its dedicated web server. Models were analyzed and visualized using UCSF ChimeraX [51] and PyMol [52].

To assess structural similarity, Root-Mean Squared Deviation (RMSD) values were determined for each pairing of the four minor pilins from *V. cholerae* to each of the four minor pilins from *S. pneumoniae* using the cealign function in PyMol [52].

Surface exposed residues were determined using the get_sasa_relative function in PyMol. A python script was used to filter through the output to identify arginine and lysine residues. Models of surface exposed arginines and lysines were generated in UCSF ChimeraX [51].

### Multiple sequence alignments

MSAs were generated using the T-Coffee server [53]. The MSAs generated using T-Coffee were then visualized using the Color Align Conservation tool in the Sequence Manipulation Suite [54]. Percentage of sequences that must agree for identity or similarity coloring to be added was set to 50%.

### Statistics

Statistical differences were assessed using GraphPad Prism software. The statistical tests used are indicated in the figure legends. Descriptive statistics for all samples, and a comprehensive list of statistical comparisons can be found in **Dataset 1**.

## ACKNOWLEDGEMENTS

This work was supported by grant R35GM128674 from the National Institutes of Health to ABD. We would like to thank Malcom Winkler, Averi McFarland, and Merrin Joseph for providing the α-MurA antibody and for helpful discussion. We would also like to thank Remi Fronzes for providing strain RL001, the parent pneumococcal strain used in this study.

---

**Fig. S1.**
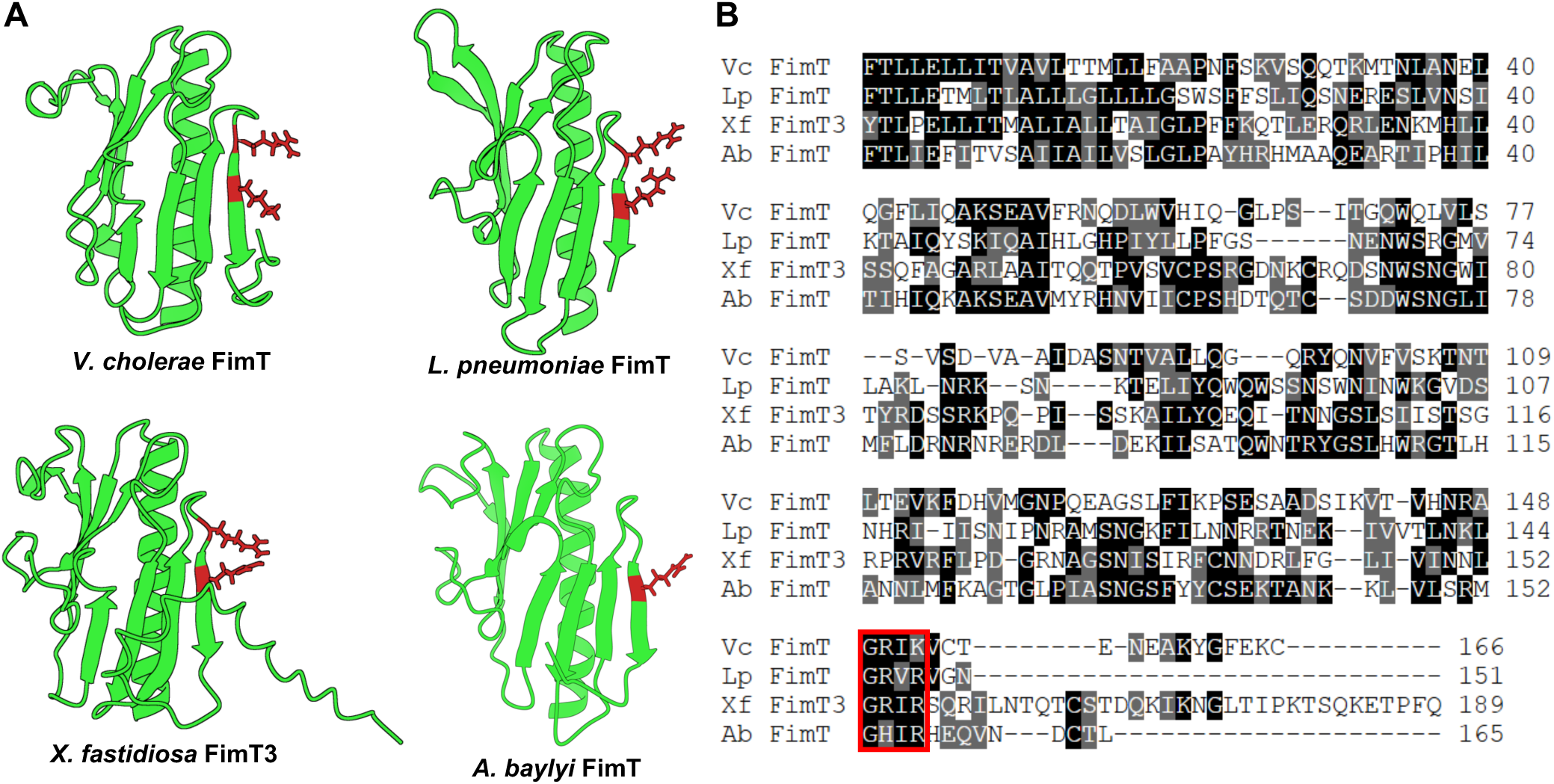
Conservation of the G-[R/K]-X-[R/K] motif in FimT homologs of diderm competence T4P. (**A**) AF- m models of the FimT homologs from the indicated bacterial species. The R/K residues from the conserved G- [R/K]-X-[R/K] motif that are required for DNA binding are highlighted in red. (**B**) Multiple sequence alignment of FimT homologs, (Vc, *V. cholerae*; Lp, *L. pneumophilla*; Xf, *X. fastidiosa*; Ab, *A. baylyi*). The G-[R/K]-X-[R/K] motif in each homolog is boxed in red. Residues that are identical are shown in black, while residues that are similar are shown in gray.

**Fig. S2.**
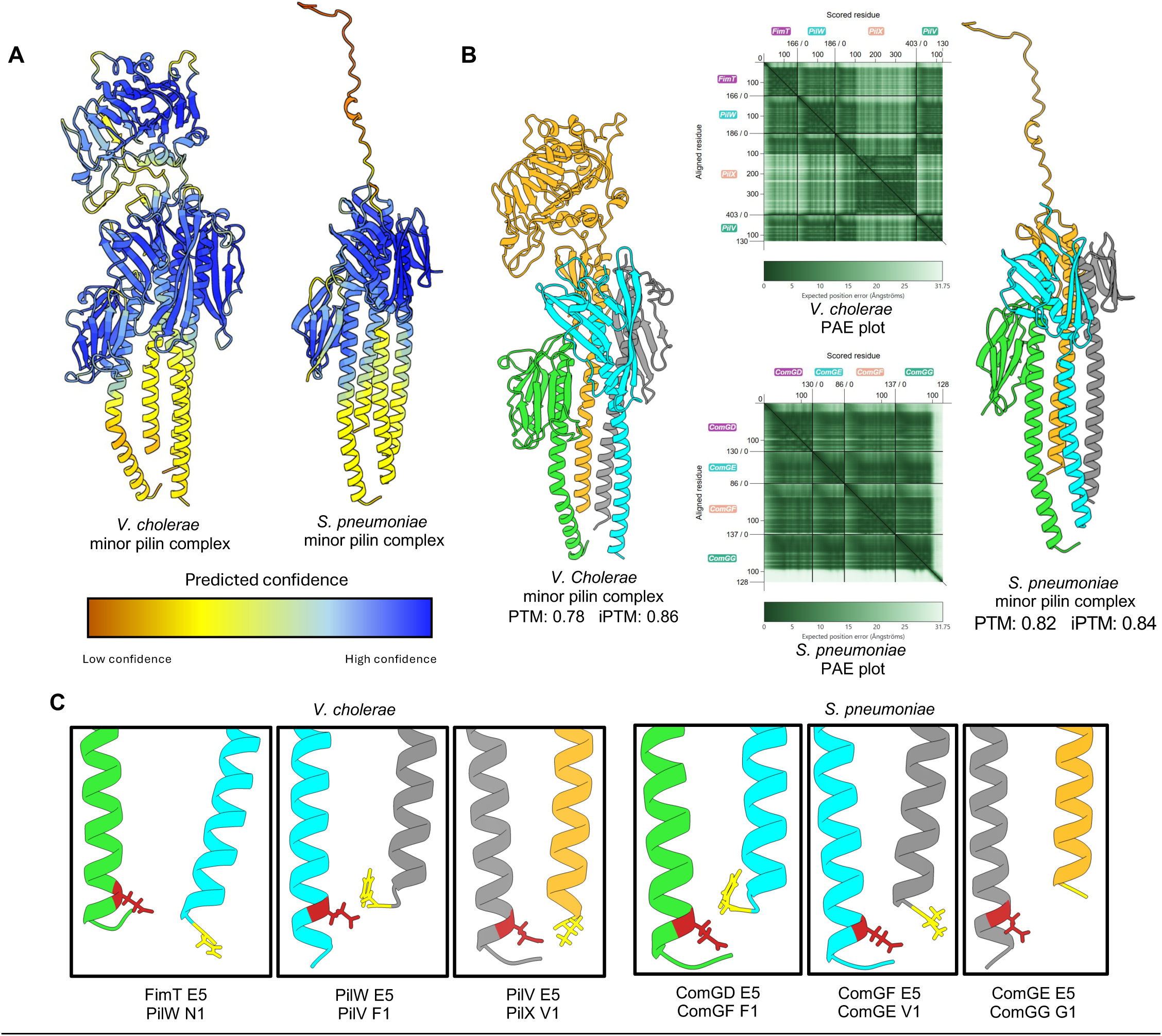
AF-m models highlight structural similarities between the competence T4P minor pilin tip complexes in *S. pneumoniae* and *V. cholerae*. (**A**) Structural predictions of competence minor pilin tip complexes from *V. cholerae* and *S. pneumoniae* colored by predicted confidence (pLDDT). (**B**) Competence minor pilin tip complexes as in (**A**) but colored by minor pilin. Also included are the pTM and ipTM scores as well as the PAE plots for each model. (**C**) Predicted E5 interactions between competence minor pilins within the complex. The E5 residue is colored red while the N-terminal residue of the neighboring pilin is colored yellow.

**Fig. S3.**
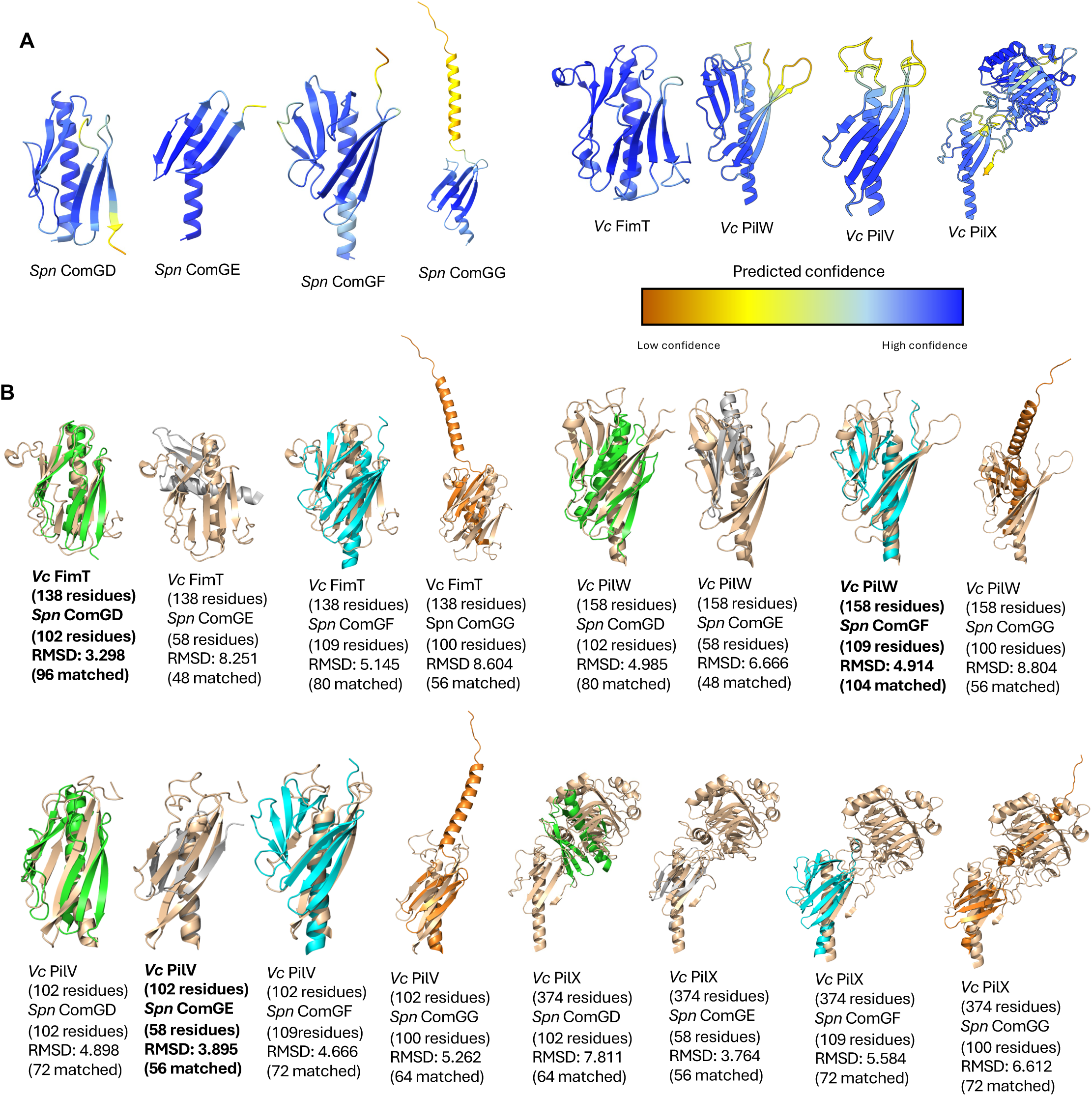
Alignment of *V. cholerae* and *S. pneumoniae* minor pilins to define structural homologs. (**A**) Individual competence minor pilins from *S. pneumoniae* (Spn) and *V. cholerae* (Vc) colored by predicted confidence score. (**B**) Alignment of the headgroups (*i.e.*, lacking residues 1-28) of the indicated minor pilins. *V. cholerae* minor pilins are shown in tan, while *S. pneumoniae* minor pilins are colored (ComGD, green; ComGE, gray; ComGF, cyan; ComGG, orange). Captions for each pairing denote the proteins aligned, the number of residues in each minor pilin, the calculated RMSD value, and the number of residues matched during the RMSD calculation. Pilin pairings with the lowest RMSD were deemed structural homologs (denoted in **bold**) with the exception of the PilX / ComGG pairing, which were deemed homologs based on the lack of an E5 (see text and **Fig. S2** for details)

**Fig. S4.**
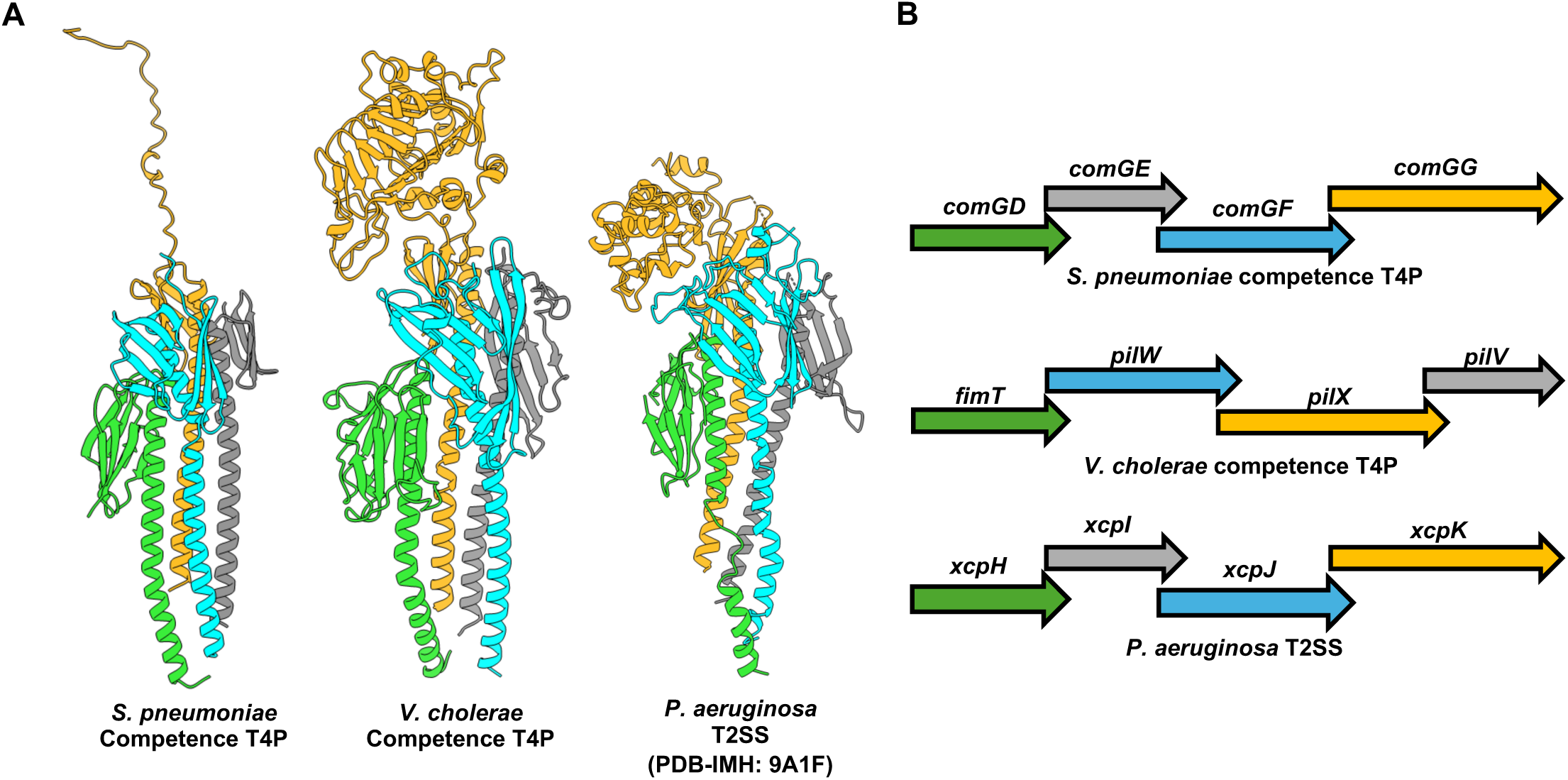
Minor pilins from monoderm competence T4P, diderm competence T4P, and T2SS share a conserved arrangement within the tip complex despite a lack of conservation in gene architecture. (**A**) AF-m models of the indicated minor pilin tip complexes and the integrative structural model of the *P. aeruginosa* T2SS minor pilin complex. (**B**) Schematic of the minor pilin operons for the indicated systems. Proteins in **A** and gene designations in **B** are color matched for ease of comparison.

**Fig. S5.**
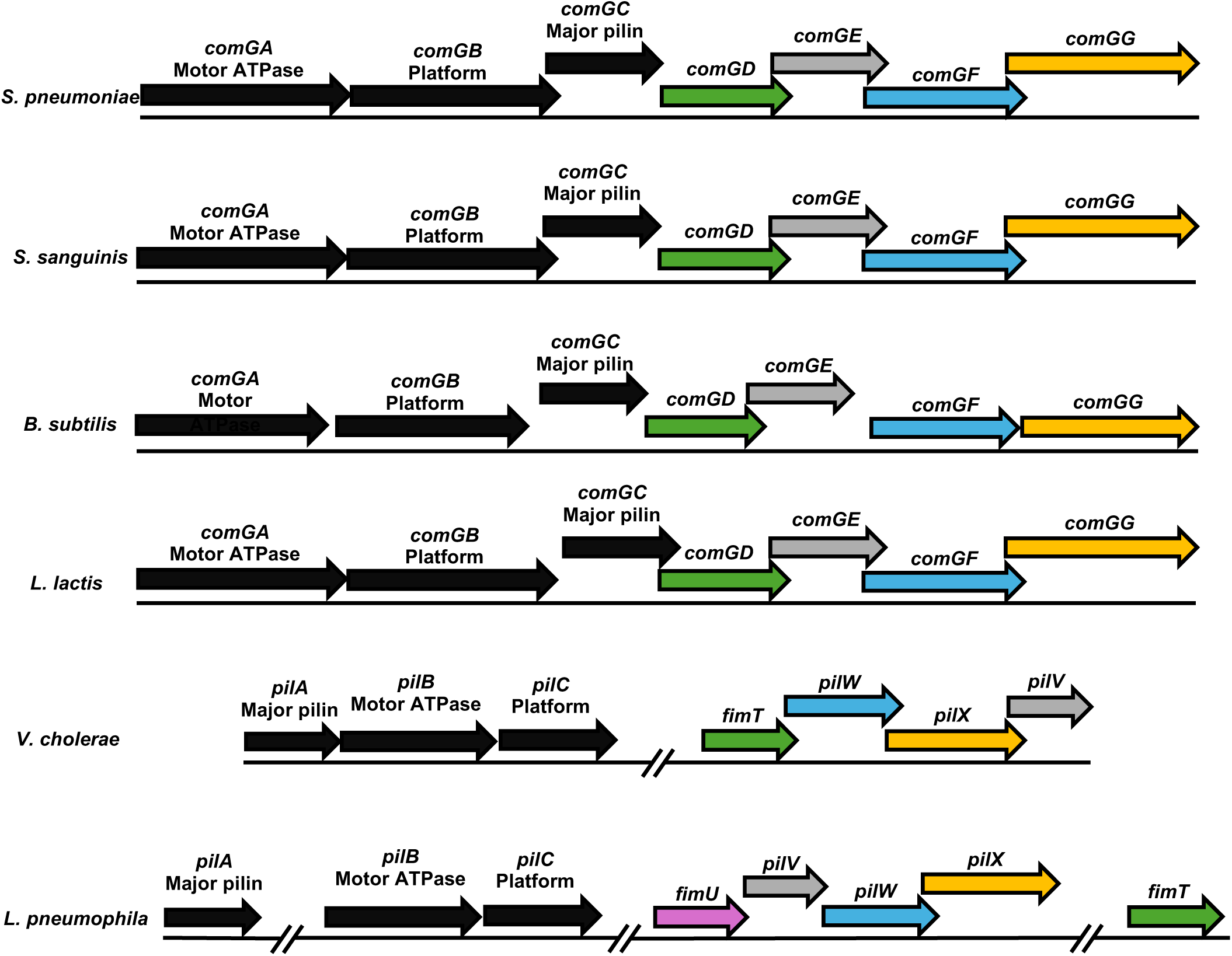
Competence T4P gene arrangements in select monoderms and diderms. Gene arrangements of the minor pilins, major pilin, motor ATPase, and platform proteins in select monoderms and diderms.

**Fig. S6.**
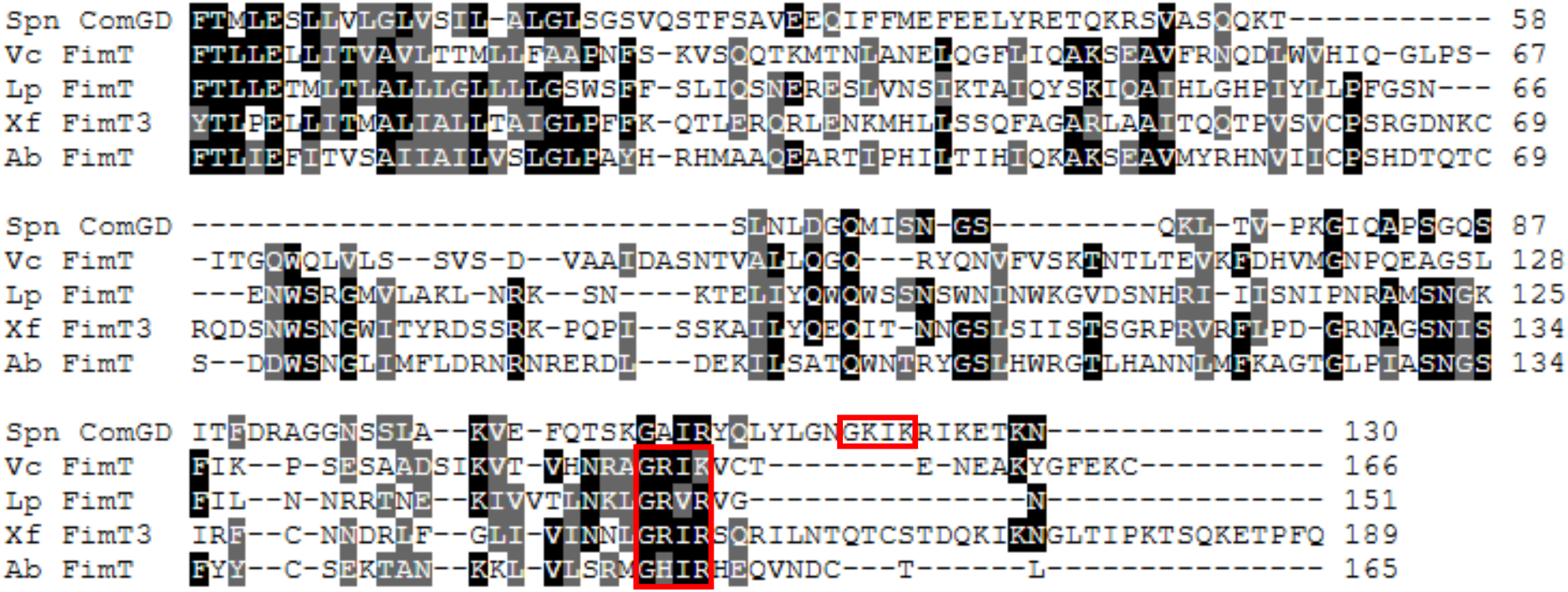
The conserved G-[R/K]-X-[G/K] motif in *S. pneumoniae* FimT is not easily identified via a sequence alignment. MSA of FimT homologs, (Spn, *S. pneumoniae*; Vc, *V. cholerae*; Lp, *L. pneumophilla*; Xf, *X. fastidiosa*; Ab, *A. baylyi*). The G-[R/K]-X-[R/K] motif in each homolog is boxed in red. Residues that are identical are shown in black, while residues that are similar are shown in gray.

**Fig. S7.**
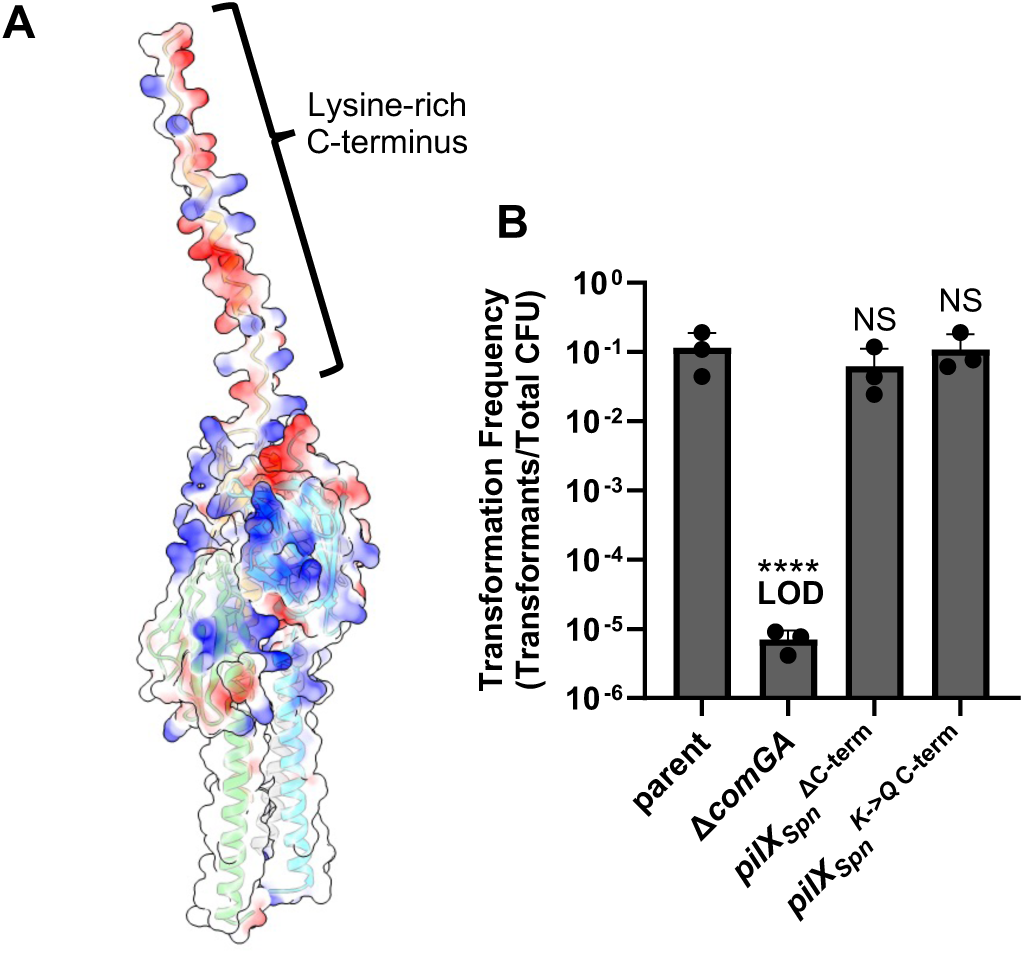
**The PilX_Spn_ C-terminus is dispensable for NT in *S. pneumoniae*.** (**A**) Electrostatic surface map of the *S. pneumoniae* minor pilin tip complex highlighting the lysine-rich C-terminus. (**B**) NT assay of the indicated *S. pneumoniae* strains. In *pilX_Spn_^ΔC-term^,* residues 101-137 were deleted. In *pilX_Spn_^K->Q C-term^*, all lysine residues within the C-terminus (residues 101-137) were mutated to glutamine (*i.e.*, V**<u>K</u>**<u>I**K**</u>EE**<u>K</u>**RD**<u>KK</u>**EEVATDSSE**<u>K</u>**VE**<u>KKK</u>**SEE**<u>K</u>**PE**<u>KK</u>**ENS was mutated to V**<u>Q</u>**<u>I**Q**</u>EE**<u>Q</u>**RD**<u>QQ</u>**EEVATDSSE**<u>Q</u>**VE**<u>QQQ</u>**SEE**<u>Q</u>**PE**<u>QQ</u>**ENS). Data in **B** is from at least 3 independent biological replicates and shown as the mean ± SD. Statistical comparisons were made by one-way ANOVA with Turkey’s multiple comparison test of the log-transformed data. NS, no significance; **** = *p* < 0.0001. LOD, limit of detection. Statistical identifiers directly above bars represent comparisons to the parent.

**Fig. S8.**
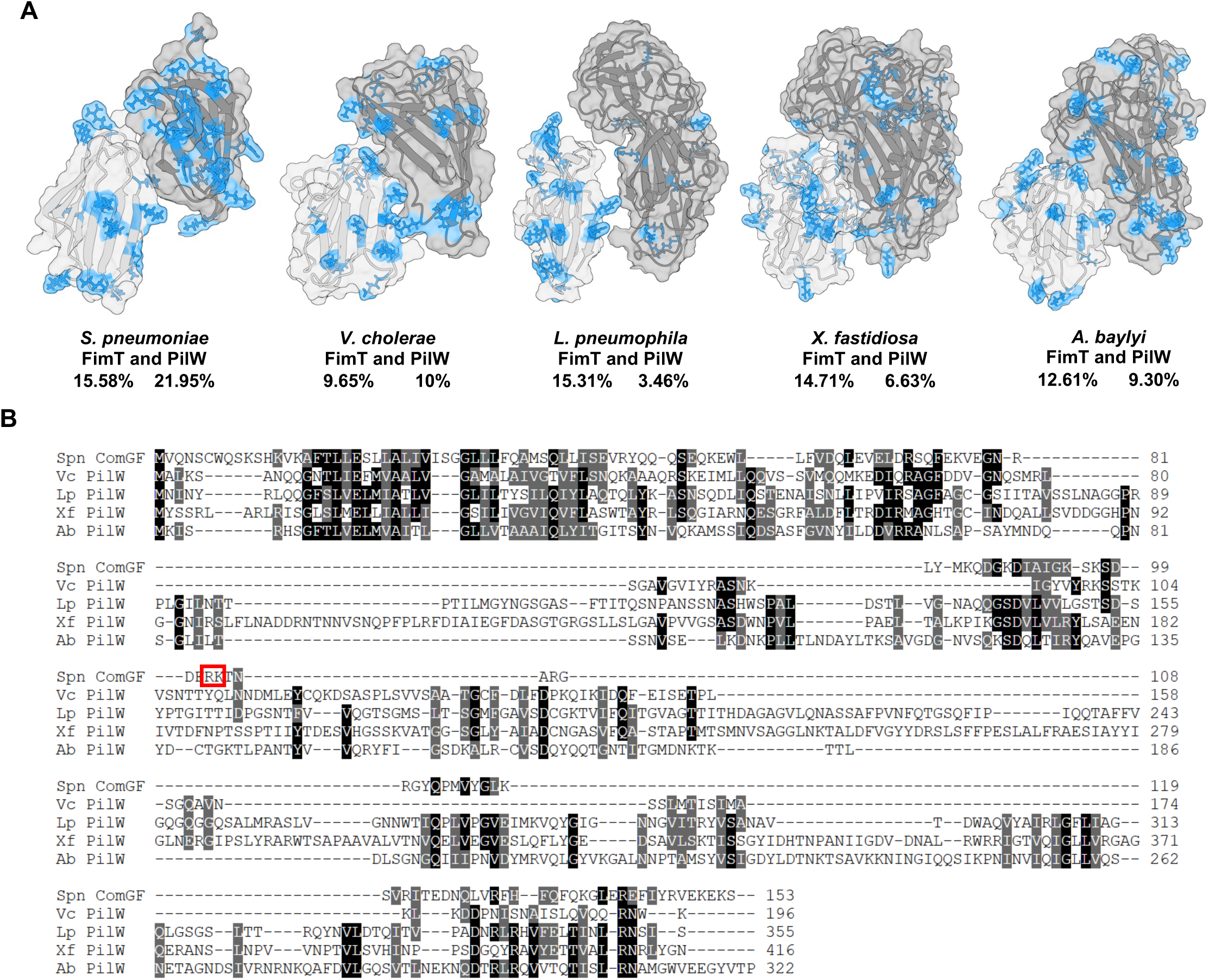
PilW_Spn_ has more surface exposed electropositive residues compared to PilW homologs in diderms. (**A**) Surface maps of FimT (light grey) and PilW (dark grey) homologs with arginine and lysine residues colored blue. The percent of surface residues that are arginines/lysines is indicated for each homolog. (**B**) MSAs of PilW homologs (Spn, *S. pneumoniae*; Vc, *V. cholerae*; Lp, *L. pneumophilla*; Xf, *X. fastidiosa*; Ab, *A. baylyi*). The residues shown to be important for DNA binding in *S. pneumoniae* are boxed in red.

**Fig. S9.**
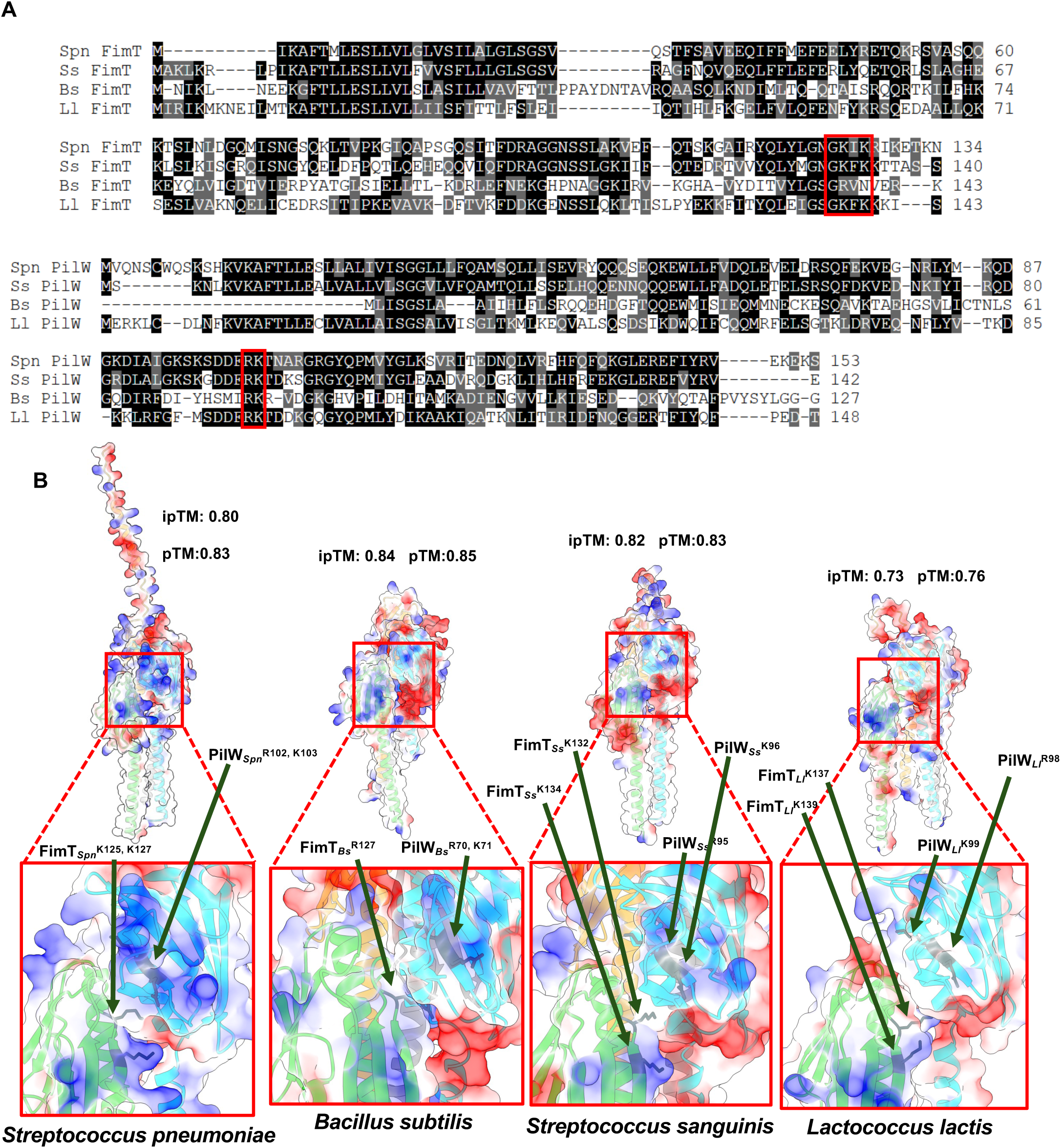
The FimT/PilW residues required for DNA-binding in *S. pneumoniae* are conserved in other monoderms. (**A**) MSAs of FimT (*i.e.*, ComGD) and PilW (*i.e.*, ComGF) homologs from four naturally competent monoderms. Spn, *S. pneumoniae;* Ss, *S. sanguinis*; Bs, *B. subtilis*; Ll, *L. lactis*. Red boxes indicate conserved residues shown to be important for NT and DNA binding in *S. pneumoniae*. Residues that are identical are shown in black, while residues that are similar are shown in gray. (**B**) Electrostatic surface maps of AlphaFold3 models of the indicated minor pilin tip complexes highlight the conserved positively-charged patch spanning FimT and PilW. Insets further highlight the positional conservation of the R/K residues shown to be critical for DNA-binding in FimTSpn and PilWSpn.

**Table S1.** Strains used in this study.

| Strain # | Genotype | Figures | Use in manuscript |
| --- | --- | --- | --- |
| TND0905/<br>SAD2468 | *P<sub>tac</sub>-tfoX*, $\Delta luxO$, *lacZ::lacI<sup>q</sup>*,<br>*pilA*<sup>S67C</sup>, $\Delta VC1807::Zeo^R$ | Fig. 1A | Parent for all *V. cholerae* strains in this study and the parent for *V. cholerae* natural transformation assays |
| NDC073/<br>SAD3694 | $\Delta fimT$, *P<sub>tac</sub>-tfoX*, $\Delta luxO$,<br>$\Delta lacZ::Spec^R$, *pilA*<sup>S67C</sup>,<br>$\Delta VC1807::Kan^R$ | Fig. 1A | $\Delta fimT$ for *V. cholerae* natural transformation assays |
| NDC0429/<br>SAD3695 | *fimT*<sup>R154Q</sup>, *P<sub>tac</sub>-tfoX*, $\Delta luxO$,<br>*lacZ::lacI<sup>q</sup>*, *pilA*<sup>S67C</sup>,<br>$\Delta VC1807::Spec^R$ | Fig. 1A | *fimT*<sup>R154Q</sup> for *V. cholerae* natural transformation assays |
| NDC0413/<br>SAD3696 | *fimT*<sup>K156Q</sup>, *P<sub>tac</sub>-tfoX*, $\Delta luxO$,<br>*lacZ::lacI<sup>q</sup>*, *pilA*<sup>S67C</sup>,<br>$\Delta VC1807::Spec^R$ | Fig. 1A | *fimT*<sup>K156Q</sup> for *V. cholerae* natural transformation assays |
| TND1035/<br>SAD2469 | $\Delta pilT::Tm^R$, *P<sub>tac</sub>-tfoX*, $\Delta luxO$,<br>*lacZ::lacI<sup>q</sup>*, *pilA*<sup>S67C</sup>, $\Delta VC1807::Zeo^R$ | Fig. 1B, C | Parent for *V. cholerae* DNA-binding assay and surface piliation microscopy |
| NDC074/<br>SAD3697 | $\Delta fimT$, $\Delta pilT::Tm^R$, *P<sub>tac</sub>-tfoX*, $\Delta luxO$,<br>$\Delta lacZ::Spec^R$, *pilA*<sup>S67C</sup>,<br>$\Delta VC1807::Kan^R$ | Fig. 1B, C | $\Delta fimT$ for *V. cholerae* DNA-binding assay and surface piliation microscopy |
| NDC0453/<br>SAD3698 | *fimT*<sup>R154Q</sup>, $\Delta pilT::Tm^R$, *P<sub>tac</sub>-tfoX*,<br>$\Delta luxO$, $\Delta lacZ::Spec^R$, *pilA*<sup>S67C</sup>,<br>$\Delta VC1807::Kan^R$ | Fig. 1B, C | *fimT*<sup>R154Q</sup> for *V. cholerae* DNA-binding assay and surface piliation microscopy |
| NDC0441/<br>SAD3699 | *fimT*<sup>K156Q</sup>, $\Delta pilT::Tm^R$, *P<sub>tac</sub>-tfoX*,<br>$\Delta luxO$, $\Delta lacZ::Spec^R$, *pilA*<sup>S67C</sup>,<br>$\Delta VC1807::Kan^R$ | Fig. 1B, C | *fimT*<sup>K156Q</sup> for *V. cholerae* DNA-binding assay and surface piliation microscopy |
| NDC0287/<br>SAD3474 | *P<sub>cilA</sub>-comGC-FLAG* downstream of<br>treR *Kan<sup>R</sup>*, $\Delta comC$ | Fig. 2A, 3B-C, 4, S3 | Strain RL001. Parent of all *S. pneumoniae* strains in this study and the parent for all *Spn* transformation assays, western blot analysis, and DNA-binding assay |
| NDC0326/<br>SAD3700 | $\Delta comGA::Erm^R$, *P<sub>cilA</sub>-comGC-FLAG*<br>downstream of treR *Kan<sup>R</sup>*, $\Delta comC$ | Fig. 2A, 3B-C, 4, S3, S4 | $\Delta comGA$ for all *Spn* transformation assays, western blot analysis, and DNA-binding assays |
| NDC0449/<br>SAD3701 | $\Delta fimT_{Spn}$, *P<sub>cilA</sub>-comGC-FLAG*<br>downstream of treR *Kan<sup>R</sup>*, $\Delta comC$ | Fig. 2A, 3B-C | $\Delta fimT_{Spn}$ for all *Spn* transformation assays, and western blot analysis |
| NDC0456/<br>SAD3702 | *fimT*<sub>Spn</sub><sup>K125Q</sup>, *P<sub>cilA</sub>-comGC-FLAG*<br>downstream of treR *Kan<sup>R</sup>*, $\Delta comC$ | Fig. 2A | *fimT*<sub>Spn</sub><sup>K125Q</sup> for *Spn* transformation assays |
| NDC0457/<br>SAD3703 | *fimT*<sub>Spn</sub><sup>K127Q</sup>, *P<sub>cilA</sub>-comGC-FLAG*<br>downstream of treR *Kan<sup>R</sup>*, $\Delta comC$ | Fig. 2A | *fimT*<sub>Spn</sub><sup>K127Q</sup> for *Spn* transformation assays |
| NDC0431/<br>SAD3704 | *fimT*<sub>Spn</sub><sup>K125Q, K127Q</sup>, *P<sub>cilA</sub>-comGC-FLAG*<br>downstream of treR *Kan<sup>R</sup>*, $\Delta comC$ | Fig. 2A, 3B-C | *fimT*<sub>Spn</sub><sup>K125Q, K127Q</sup> for *Spn* transformation assays and western blots |
| NDC0432/<br>SAD3705 | *pilW*<sub>Spn</sub><sup>R102Q</sup>, *P<sub>cilA</sub>-comGC-FLAG*<br>downstream of treR *Kan<sup>R</sup>*, $\Delta comC$ | Fig. 3B-C | *pilW*<sub>Spn</sub><sup>R102Q</sup> for *Spn* transformation assays and western blots |
| NDC0443/<br>SAD3706 | *fimT*<sub>Spn</sub><sup>K125Q, K127Q</sup>, *pilW*<sub>Spn</sub><sup>R102Q</sup>,<br>*P<sub>cilA</sub>-comGC-FLAG* downstream of<br>treR *Kan<sup>R</sup>*, $\Delta comC$ | Fig. 3B | *fimT*<sub>Spn</sub><sup>K125Q, K127Q</sup>, *pilW*<sub>Spn</sub><sup>R102Q</sup> for *Spn* transformation assays |
| NDC0466/<br>SAD3708 | *fimT*<sub>Spn</sub><sup>R116Q</sup>, *P<sub>cilA</sub>-comGC-FLAG*<br>downstream of treR *Kan<sup>R</sup>*, $\Delta comC$ | Fig. 3B | *fimT*<sub>Spn</sub><sup>R116Q</sup> for *Spn* transformation assays |
| NDC0483/<br>SAD3708 | $\Delta pilW_{Spn}$, *P<sub>cilA</sub>-comGC-FLAG*<br>downstream of treR *Kan<sup>R</sup>*, $\Delta comC$ | Fig. 3B-C | $\Delta pilW_{Spn}$ for *Spn* transformation assays, and western blots |
| NDC0484/<br>SAD3709 | *pilW*<sub>Spn</sub><sup>R102Q, K103Q</sup>, *P<sub>cilA</sub>-comGC-FLAG*<br>downstream of treR *Kan<sup>R</sup>*, $\Delta comC$ | Fig. 3B-C | *pilW*<sub>Spn</sub><sup>R102Q, K103Q</sup> for *Spn* transformation assays and western blots |
| NDC0486/<br>SAD3710 | *pilW*<sub>Spn</sub><sup>K103Q</sup>, *P<sub>cilA</sub>-comGC-FLAG*<br>downstream of treR *Kan<sup>R</sup>*, $\Delta comC$ | Fig. 3B-C | *pilW*<sub>Spn</sub><sup>K103Q</sup> for *Spn* transformation assays and western blots |
| NDC0487/<br>SAD3711 | *fimT*<sub>Spn</sub><sup>K105Q</sup>, *P<sub>cilA</sub>-comGC-FLAG*<br>downstream of treR *Kan<sup>R</sup>*, $\Delta comC$ | Fig. 3B | *fimT*<sub>Spn</sub><sup>K105Q</sup> for *Spn* transformation assays |
| NDC0488/<br>SAD3712 | *pilW*<sub>Spn</sub><sup>K107Q</sup>, *P<sub>cilA</sub>-comGC-FLAG*<br>downstream of treR *Kan<sup>R</sup>*, $\Delta comC$ | Fig. 3B | *pilW*<sub>Spn</sub><sup>K107Q</sup> for *Spn* transformation assays |
| NDC0489/<br>SAD3713 | *pilW<sub>Spn</sub><sup>K109Q</sup>*, *P<sub>cilA</sub>-comGC-FLAG*<br>downstream of *treR Kan<sup>R</sup>, ΔcomC* | Fig. 3B | *pilW<sub>Spn</sub><sup>K109Q</sup>* for *Spn* transformation assays |
| NDC0490/<br>SAD3714 | *fimT<sub>Spn</sub><sup>K125Q, K127Q</sup>*, *pilW<sub>Spn</sub><sup>K103Q</sup>*, *P<sub>cilA</sub>-*<br>*comGC-FLAG* downstream of *treR Kan<sup>R</sup>, ΔcomC* | Fig. 3B | *fimT<sub>Spn</sub><sup>K125Q, K127Q</sup>*, *pilW<sub>Spn</sub><sup>K103Q</sup>* for *Spn*<br>transformation assays |
| NDC0491/<br>SAD3715 | *fimT<sub>Spn</sub><sup>K125Q, K127Q</sup>*, *pilW<sub>Spn</sub><sup>R102Q, K103Q</sup>*, *P<sub>cilA</sub>-comGC-FLAG*<br>downstream of *treR Kan<sup>R</sup>, ΔcomC* | Fig. 3B-C,<br>Fig 4, S3 | *fimT<sub>Spn</sub><sup>K125Q, K127Q</sup>*, *pilW<sub>Spn</sub><sup>R102Q, K103Q</sup>* for *Spn*<br>transformation assays, western blots, and DNA-binding assays |
| NDC0430/<br>SAD3716 | *comGC<sup>S66C</sup>*, *P<sub>cilA</sub>-comGC-FLAG*<br>downstream of *treR Kan<sup>R</sup>, ΔcomC* | Fig. S4 | Parent for *Spn comGG* transformation assays |
| NDC0494/<br>SAD3717 | *comGG<sup>Δ101-137</sup>*, *comGC<sup>S66C</sup>*, *P<sub>cilA</sub>-*<br>*comGC-FLAG* downstream of *treR Kan<sup>R</sup>, ΔcomC* | Fig. S4 | *comGG<sup>ΔC-term</sup>* for *Spn* transformation assays |
| NDC0495/<br>SAD3718 | *comGG<sup>K->Q</sup>*, *comGC<sup>S66C</sup>*, *P<sub>cilA</sub>-*<br>*comGC-FLAG* downstream of *treR Kan<sup>R</sup>, ΔcomC* | Fig. S4 | *comGG* C-term K->Q for *Spn* transformation<br>assays |

\*Double identifiers under “Strain #” refers to the same strain that has been stocked in two independent strain collections.

**Table S2.**
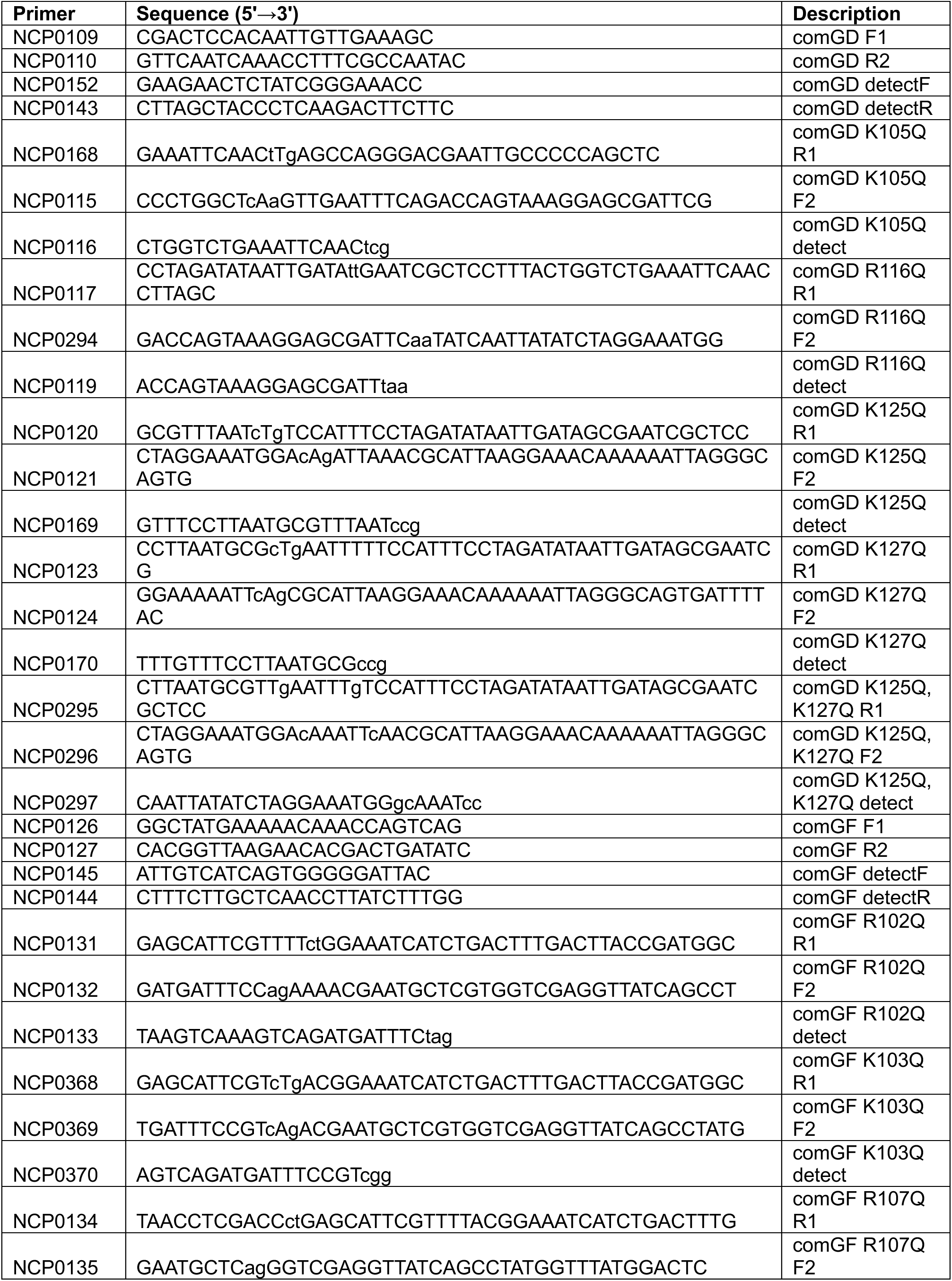
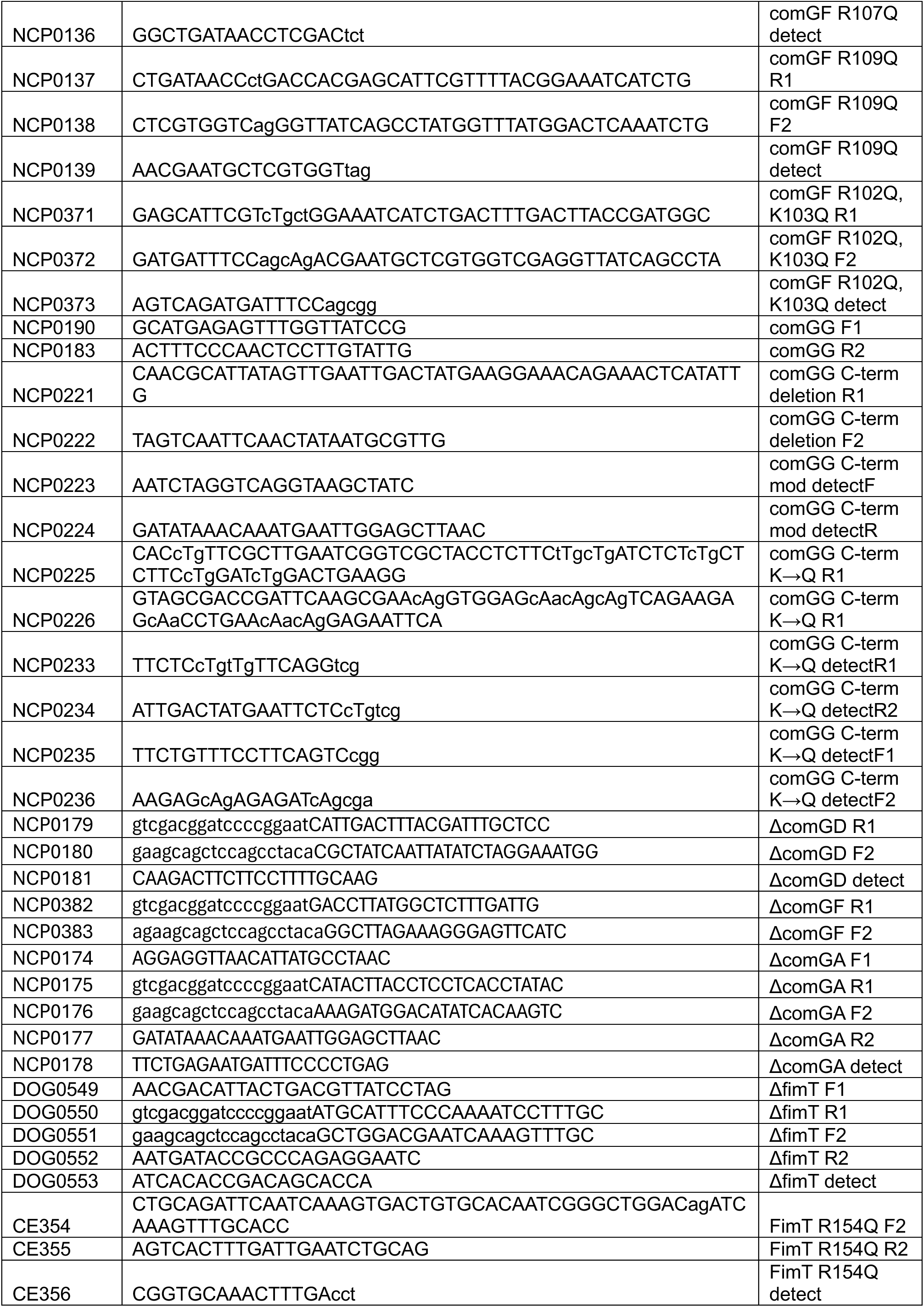
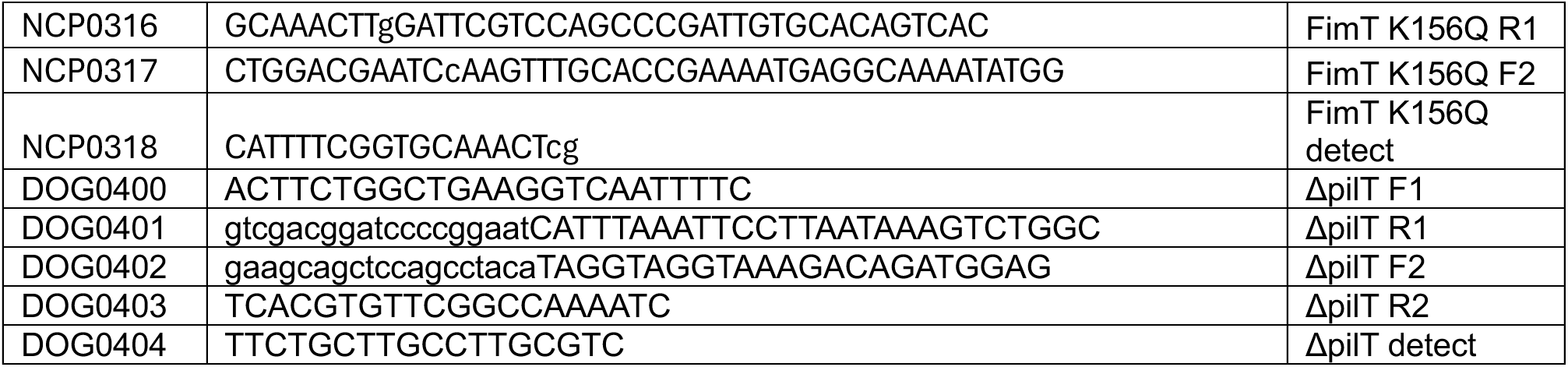
Primers used in this study.

